# Refining Feulgen: low-cost and accurate genome size measurements for everyone

**DOI:** 10.1101/2025.08.29.673164

**Authors:** Mohammed M.Tawfeeq, Ursula Swaelus, Florence Rodriguez Gaudray, Jenny Arrensdorff, Felipe Ennes Silva, Tristan Permentier, Laurent Grumiau, Thomas Verdebout, Jean-François Flot

## Abstract

Feulgen image analysis densitometry (FIAD) has been extensively used for decades to measure genome sizes, but is usually considered less precise and less reproducible compared to other methods such as flow cytometry (FCM). However, FCM requires fresh tissue samples, which is a major impediment to using it on specimens collected in the field far from the lab. By contrast, FIAD works on ethanol-preserved samples kept at room temperature, which is a huge advantage for biodiversity genomics. In this study, we present refinements on the Feulgen protocol and downstream analyses that produce measurements at least as reliable and precise as those obtained using whole-genome sequencing. We verify using seven standards (including both animals and plants) that our refined approach yields accurate results across a 440-fold range of genome sizes. To illustrate the usefulness of our approach, we use it to measure for the first time the genome size of the red bald uakari monkey *Cacajao rubicundus* using the American cockroach *Periplaneta americana* and the black garden ant *Lasius niger* as standards, and confirm our 2.67 pg result by comparing it with a whole-genome assembly obtained from nanopore sequencing of the same individual.

## 1 Introduction

Measuring genome size is a critical prerequisite for modern biological research, underpinning everything from strategic planning of genome sequencing projects to fundamental investigations of genome evolution. Genome sizes in eukaryotes are commonly reported as C-values, defined as half of the DNA content of a diploid somatic nucleus [1] and expressed in (pg) of DNA or in (Mb) or (Gb) of bases (1 pg = 978 Mb). C-values have profound biological implications: for instance, a larger genome size has been found to be positively correlated with an increased likelihood of extinction in both plants [2, 3] and vertebrates [4]. This has spurred the creation of comprehensive online databases to systematically catalog C-value data across various species [5].

The current state-of-the-art for measuring genome size is flow cytometry [6]. This technique is widely adopted for its speed and precision, providing reliable measurements with clear estimates of confidence. However, its utility is significantly hampered by practical constraints. The method is costly in terms of both equipment and reagents, and it requires large numbers of intact nuclei [7], which in many cases can only be obtained from fresh tissues. These requirements pose major logistical challenges for large-scale studies, particularly those involving samples collected from remote field locations. Other frequently used techniques, such as qPCR [8, 9], k-mer-based approaches [10] and genome assembly, also present drawbacks, such as the need for species-specific optimization for qPCR, lack of precision of k-mer based approaches, and assembler-specific variations in behavior when dealing with heterozygous or otherwise polymorphic regions [11].

Historically, one of the foundational methods for DNA quantification has been the Feulgen reaction, which was first proposed to stain DNA in cells and tissues by Robert Feulgen and Heinrich over a century ago [12] and later used to measure genome sizes [13]. The key advantage of Feulgen image analysis densitometry (FIAD) for genome size measurement is that it can be used with ethanol-preserved specimens, making it ideal for surveying specimens from field collections. However, FIAD is often considered as less precise and less quantitative than flow cytometry [14].

In the present study, we refined the classic FIAD approach to transform it into a cost-effective and accurate method, well-suited for surveying genome sizes in large-scale studies. We used our optimized protocol to produce a robust genome size estimate for the red-faced uakari monkey (*Cacajao rubicundus*, a primate species endemic to the Brazilian Amazon Rainforest, with a distribution restricted to the middle Solimões River basin [15]), that we subsequently validated using whole-genome sequencing and assembly.

## 2 Materials and Methods

Conducting Feulgen protocol in previous experiments presented inconsistencies ranging from the chemicals’ preparation to the software and camera quality, leading to low reproducibility. To address these issues, we developed a robust, optimized workflow that is broadly applicable across diverse animal and plant tissues. The key differences in our refined method are the use of multiple reference standards and the introduction of a novel genome size histogram. Previous protocols relied on a single standard, a critical limitation that precludes the verification of staining efficacy to confirm whether the staining worked as expected. By integrating several standards, our approach introduces essential diagnostic quality controls that enable users to validate the staining efficacy and confirm assay overall reliability before proceeding with data analysis. The detailed steps of this optimized workflow, supplemented by comprehensive troubleshooting guidelines, are described below.

### 2.1 Sample Collection

To test the method across a broad dynamic range, we selected a panel of seven reference species with well-characterized C-values. The animal standards are conveniently reared at the Department of Organismal Biology of the Université libre de Bruxelles (ULB) for teaching and research purposes: the bdelloid rotifer *Adineta vaga* (C-value = 0.10 pg ~ 100.8 Mb [16]), the black garden ant, *Lasius niger* (C-value = 0.30 pg ~ 293.4 Mb [17]), and the American cockroach, *Periplaneta americana* (C-value = 3.41 pg ~ 3,334 Mb) [18, 19]. We used the dissected brain of a female *P. americana* and three individual *L. niger* female worker ants (after removing their abdomens). The selection of female ant workers ensured the analysis of diploid somatic cells, as opposed to the haploid cells of male ants. Using brain tissues (for the cockroach) and avoiding abdomens (for the ant) was in order to avoid actively dividing tissues (such as the gut) that contain a mix of G1 and G2 cells.

Plant standards included: *Agave americana* (C-value = 7.95 pg), *Clivia miniata* (C-value = 19 pg), *Haemanthus albiflos* (C-value = 36 pg), and *Scadoxus multiflorus* (C-value = 44.2 pg) [20, 21], with leaf tissues obtained from the Meise Botanical Garden (Meise, Belgium; IPEN numbers 19083990, 2006137226, 19200124, and 19411603, respectively).

For testing our optimized protocol, we used a muscle tissue sample to estimate the genome size of a female red bald-headed uakari, *Cacajao rubicundus*. This sample was provided by the Mamirauá Institute for Sustainable Development (permit numbers: CITES 19BR033597/DF; SISBIO 55777).

Tissues from all specimens were preserved in 96% ethanol after collection and until processing.

### 2.2 Reagents and Equipment

The essential laboratory equipment includes an incubator, a fume hood, and a digital balance. For image acquisition, a TOUPCAM™ FHD camera (TOUPTEK Photonics CO., LTD) was utilized, connected to a computer running ToupView software. Subsequent image analysis was performed using ImageJ, and statistical analyses were conducted in RStudio.

### 2.3 Slide preparation

A tissue sample, approximately 0.4 cm in length and 0.3 cm in width, was excised from the specimen using a sterilized razor blade or scissors. The excised tissue was placed directly onto a clean, dry microscope slide, which had been pre-cleaned with absolute alcohol.

Following placement on the slide, one to two drops of 40% acetic acid were added to the tissue. The tissue was then meticulously chopped into fine pieces directly on the slide with a sterilized razor blade. To prevent cell clumping and ensure a homogenous single-cell layer, additional drops of 40% acetic acid were applied, and the resulting cell suspension was carefully spread in a circular motion across the slide.

All prepared slides were left in a horizontal position at room temperature under the hood for 3 hours to allow for the complete evaporation of the acetic acid. Subsequently, the slides were fully immersed in 96% ethanol and allowed to air dry completely. Finally, the finished slides were stored in a dark, room-temperature environment for 48 hours before imaging and analysis to ensure optimal nuclei staining. The same detailed procedure was followed to prepare the standards. In practice, researchers can begin the protocol by selecting at least two reference standards that closely match the expected C-value of their target organism.

### 2.4 Fixation

This step is crucial, as it dehydrates cells while preserving the tissue’s structure in its original state, cross-linking nuclear proteins, and preventing DNA loss or degradation [14]. The fixation reagent was prepared by mixing 100% methanol, 37% formaldehyde and 90% acetic acid solutions in relative volumes (85:10:5). The prepared slides of standards and samples were placed in the slide rack then carefully immersed in a glass tank filled with the fixation reagent. The tank was sealed with parafilm and aluminum foil, then stored in a dark environment for 24 hours at 25 °C.

### 2.5 Washing with distilled water

The slide rack was carefully removed from the fixative tank and immersed in distilled water for 5 minutes. This was repeated a second time to remove residual chemicals.

### 2.6 Hydrolysis

To depurinate DNA and create free aldehyde groups [22], we immerged the slide rack in a fresh glass tank containing hydrochloric acid (HCl) at a 5M concentration. The container was sealed with parafilm, covered with aluminum foil, and stored in an incubator for 2 hours [23] at 25 °C. The rinsing procedure described in step 2.5 was then performed again.

### 2.7 Staining

Binding of the fuchsin on the free aldehyde groups formed on the DNA molecule during hydrolysis [22] was performed by placing the slide rack in a new glass tank containing Schiff reagent, then sealing the tank with parafilm and aluminum foil before incubating for 2 hours [14] in a dark setting at 25 °C.

### 2.8 Washing with sodium disulphite

A sodium disulphite solution was prepared to a volume of 200 ml by dissolving 1g of sodium disulphite in 10 ml of distilled water, adding 188 ml of distilled water, then finally adding 2 ml of HCl 5M. To remove and de-colorize unbound fuchsin stain, [14], the slides were placed in a tank filled with freshly prepared sodium disulphite buffer for 5 minutes, then a second time. Disulfite residues were then eliminated by performing two rinses in distilled water, as in section 2.5.

### 2.9 Dehydration

Finally, a few drops of 70% ethanol were added onto each processed slides and they left to dry for approximately 15 minutes until all the ethanol had evaporated. This was then repeated with 90% ethanol. The dehydrated slides were subsequently stored in a dark, dry environment till microscopy.

### 2.10 Microscopy

A few drops of immersion oil was applied onto each slide before covering them with a coverslip [14]. One or two drops of immersion oil were added on the coverslip itself, then a 100x oil-immersion objective was used to examine the slide on a KERN Optic microscope. The microscope settings, including maximum light intensity, the condenser’s distance from the mechanical stage, and the iris diaphragm aperture, were maintained fixed to ensure consistent measurements across all standards and samples. Since DNA staining gradually fades over time, we photographed both standards and samples within 3 weeks maximum after performing the Feulgen assay; photographing beyond this time window gave us inaccurate genome size estimates.

A 5-megapixel ToupCam digital color camera attached to the microscope was used to observe the slides; images were captured using the ToupView software (Windows version 4.11.19) at a resolution of 2560 × 1922 pixels.

### 2.11 Image analysis

Images were analyzed using ImageJ [24], an open-source tool for the analysis of scientific images developed by the National Institute of Health. We analyzed 30 nuclei per sample (clumped or overlapping nuclei were excluded from the analysis). Three important measures were taken: the area of the nucleus (A), the mean gray value of the nucleus (GVN), and the mean gray value of the background (GVB) (Fig. 1). From these we inferred the optical density (OD) of each nucleus as the difference between the mean gray value of the background and the mean gray value of the nucleus.

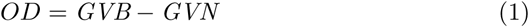

**Fig. 1.**
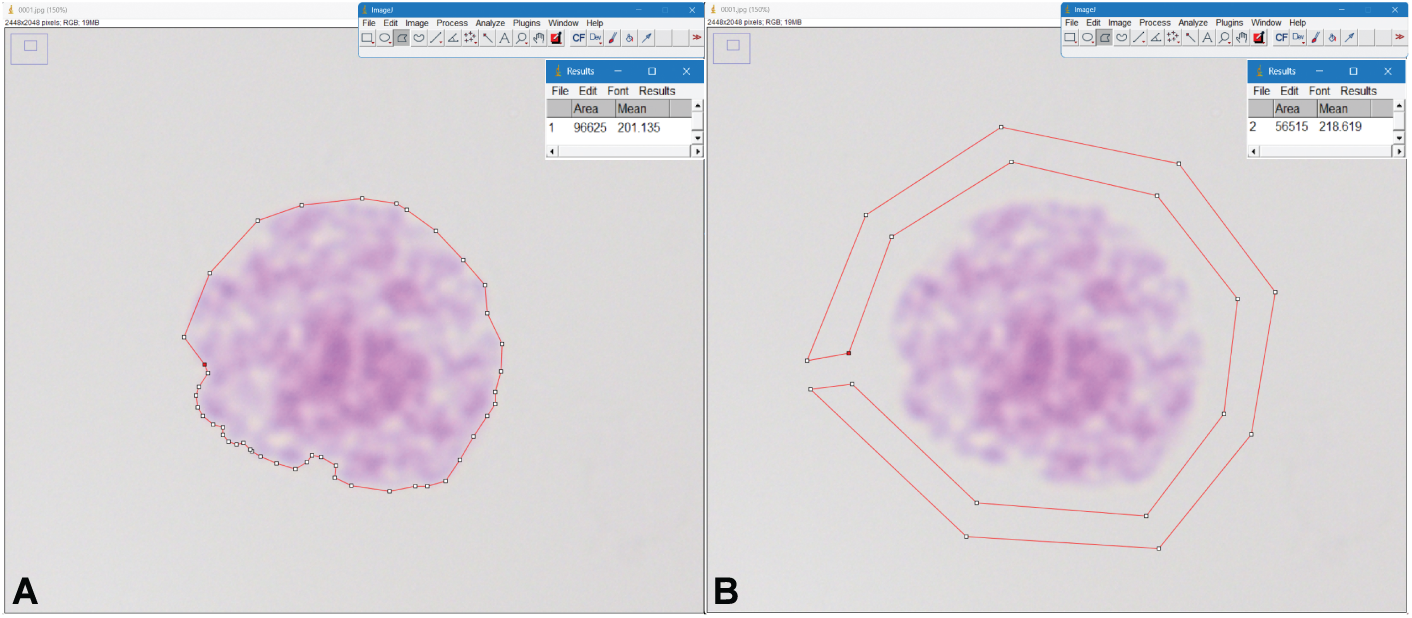
Measurements of the nuclear area size and its light absorbance intensity from the same individual nucleus. (A) A nucleus boundary is outlined using ImageJ to extract the nucleus’s area size and its mean gray value. (B) An external boundary of the background surrounding the nucleus is outlined to estimate the mean gray value of the background.

This OD value can then be multiplied by the area of the nucleus (A) to obtain the integrated optical density (IOD), which is proportional to the genome size (C-value) of the specimen under investigation.

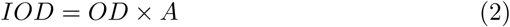

For the sake of consistency of analyses, we used only G1 nuclei for our downstream analyses and discarded G2 ones (that contain a double amount of DNA compared to G1 nuclei, resulting in an IOD approximately two times larger). In most eukaryotic species, the majority of nuclei are in G1 phase, because G1 takes longer than G2 during the phase cycle. However, there are some exceptions that users need to take into account, such as in the ”blob” *Physarum polycephalum* where almost all nuclei are in G2 [25]. In such case, users should take care to divide by two the observed 2C genome size inferred using our method, in order to recover the actual C-value. A schematic overview of the complete laboratory workflow is presented in Fig. 2.

**Fig. 2.**
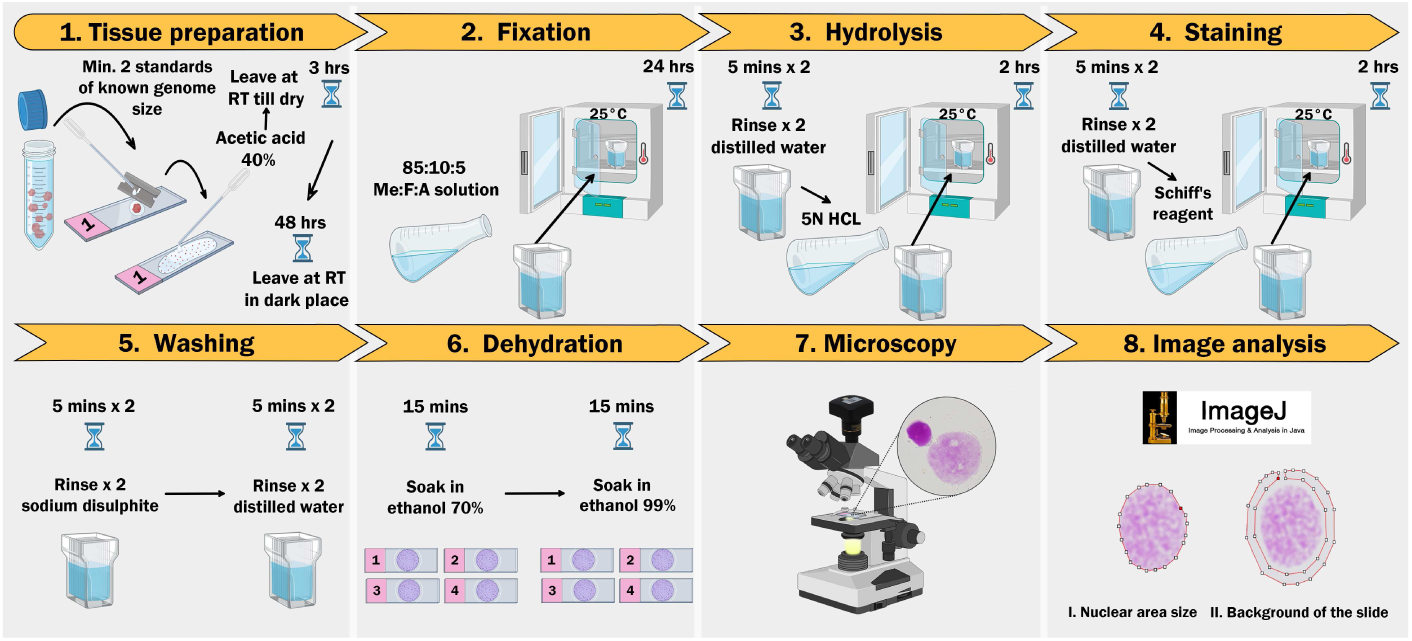
The refined Feulgen Image Analysis Densitometry protocol. Created in BioRender. M. Tawfeeq, M. (2026) https://BioRender.com/97wcy9l.

### 2.12 Data quality control

To control the quality of the measurements performed, we verified that they met the two basic assumptions of the Feulgen method. The first assumption is that nearly all nuclei of a given specimen contain the same amount of DNA, regardless of their area. This was tested by plotting 1/OD (the reverse of the optical density) vs. A (the nuclear area). The assumption is met for a given specimen if the resulting points fall on a line that passes nearby the origin of the plot, i.e., 1/OD is proportional to A, meaning that IOD = OD x A is constant for this specimen (Fig. 3).

**Fig. 3.**
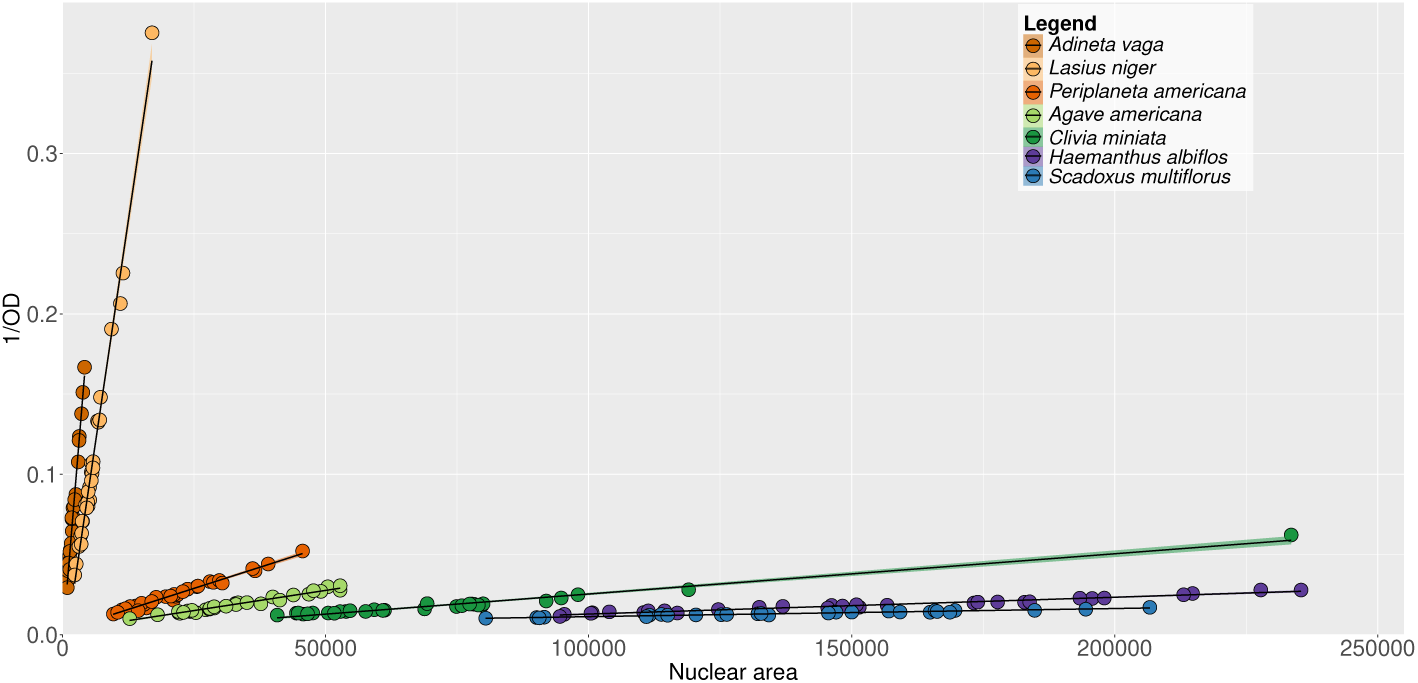
Quality control confirming that the amount of coloration in the G1 nuclei of each analysed standard is constant regardless of their area size. The shaded bands around the regression lines indicate the 95% confidence intervals.

The second assumption is that IOD is proportional to the genome size (C-value) of the species, i.e., the Schiff reagent colored DNA in a stoichiometric fashion (Fig. 5). To verify that this assumption holds across a large range of genome sizes covering most measured species, we checked that the average IODs of seven different reference standards, when plotted against their known C-values, fell on a line that passed nearby the origin of the plot, which was verified using linear regression analysis to demonstrate a highly linear relationship. The reproducibility of the method was evaluated by comparing two independent experimental runs using the same standard specimens. The two runs produced almost identical regression lines, in close agreement with the ideal proportionality expected (slope = 1) (Fig. 4). Although we used seven random standards to demonstrate the wide dynamic range of our optimized method, routine implementation of this method requires no more than two reference standards. Therefore, users can adapt the protocol by choosing any locally accessible standards that match the organism of interest and its expected genome size.

**Fig. 4.**
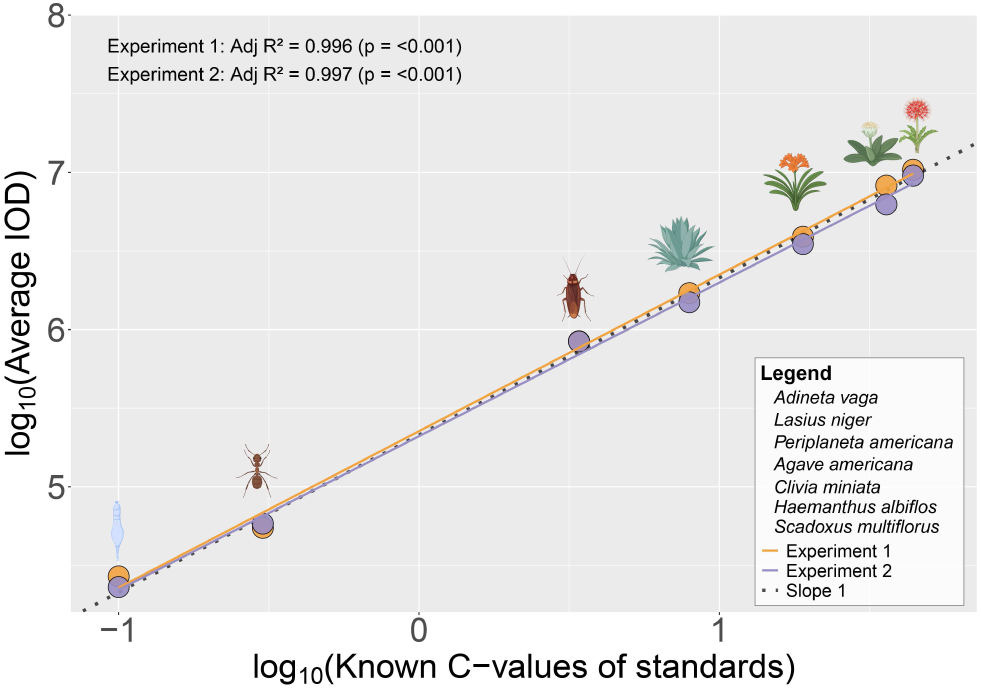
Quality control confirms that the average nuclear coloration for each standard species is proportional to its known C-value. Reproducibility is supported by the overlap of two independent experimental runs using the same standard specimens, which yielded almost identical linear regressions that closely fit with a slope of 1.0. Species pictures were created in BioRender. M.Tawfeeq, M. (2026) https://BioRender.com/0z4htw9.

**Fig. 5.**
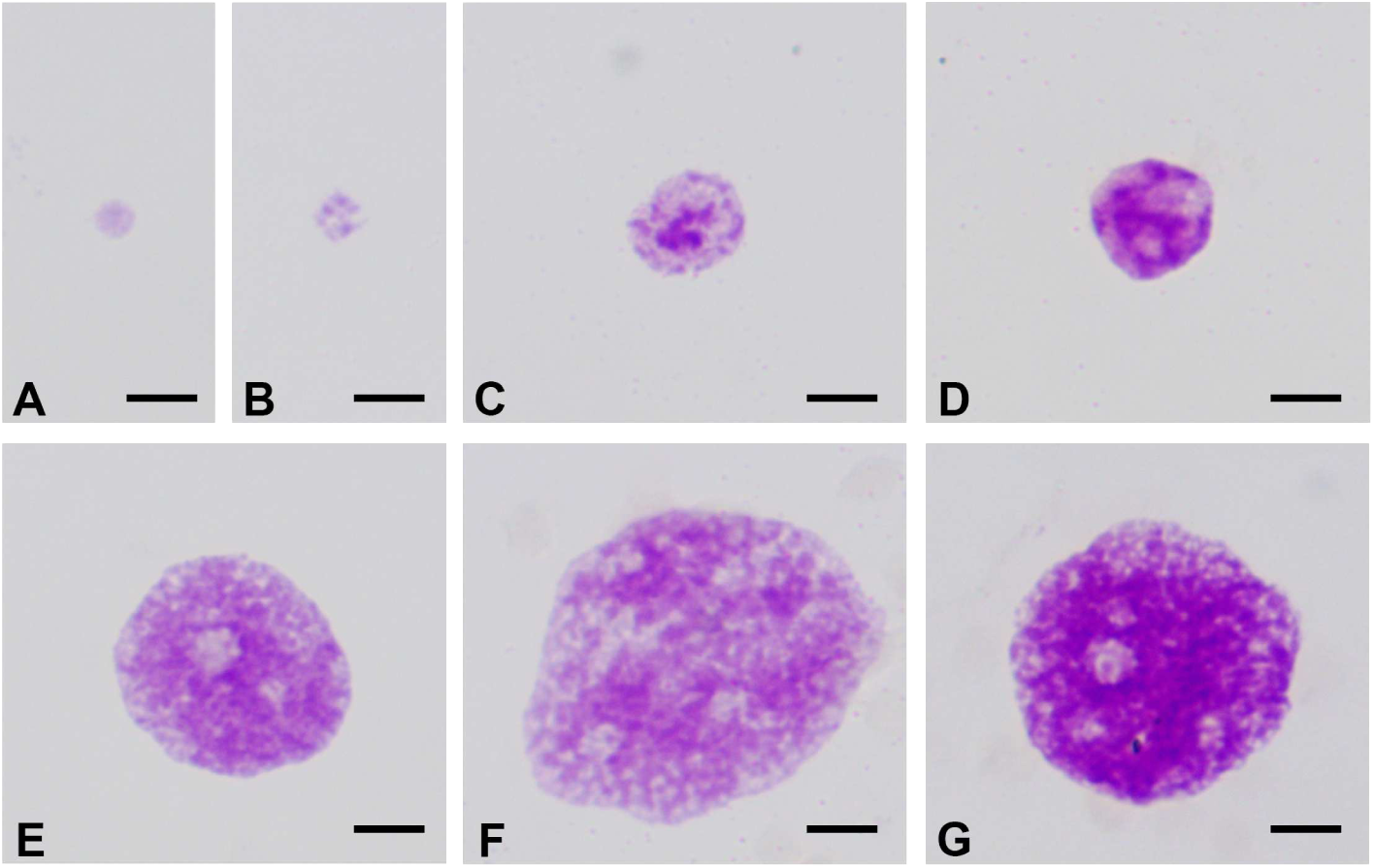
Intact nuclei from seven reference standards spanning the smallest to the largest genome sizes: (A) *Adineta vaga*, (B) *Lasius niger*, (C) *Periplaneta americana*, (D) *Agave americana*, (E) *Clivia miniata*, (E) *Haemanthus albiflos*, and (G) *Scadoxus multiflorus*. The arrangement illustrates a clear visual relationship between nuclear area, Feulgen staining intensity, and genome size from the smallest (A) to the largest genome (G). Scale bar: 5 *µ*m.

### 2.13 Genome size calculation

Once verified that data passed both of the two quality controls detailed above, the C-value of the sample (*C_sample_*) was estimated from its IOD (*IOD_sample_*). Since the assumption is that IOD is proportional to C-value, we have

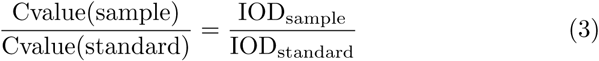

We calculated this C-value for each combination of one nucleus of the sample and one nucleus of the one of the standard: as there were 30 nuclei for our *C. rubicunduns* sample and 30 nuclei for each of our two standards, this resulted in a set of 1800 semi-independent estimates of the C-value of *C. rubicunduns*. These values were then visualized as a histogram superposed with the corresponding Gaussian kernel density, the mode of which yielded our final estimate of C-value of *C. rubicunduns* (expressed in pg of DNA) (Fig. 7). To estimate its standard deviation (*σ*), we measured the width of the Gaussian kernel density distribution at 0.882 time the height of the mode [26]. The coefficient of variation (*CV_sample_*) was then calculated as the ratio of this standard deviation (*σ_sample_*) to the modal C-value of the generated data matrix and expressed as a percentage (Fig. 7).

### 2.14 DNA isolation, sequencing, and data processing

Genomic DNA was extracted from the muscle tissue of the same *C. rubicundus* specimen as for Feulgen, using Qiagen’s MagAttract HMW DNA Kit (Ref: 67563). A ligation sequencing SQK-LSK114 Nanopore library was prepared according to the manufacturer’s protocol and sequenced on a FLO-PRO114M flow cell with R10.4.1 pore proteins on a PromethION P2-solo machine (Oxford Nanopore Technologies, Oxford, UK). Sequencing was performed at the Evolutionary Biology and Ecology research unit, ULB (Brussels, Belgium). Nanopore base calling was performed using Guppy v3.2.10. The resulting reads were assembled de novo using Flye [27]. The quality and completeness of the assembly were assessed and calculated using K-mer Analysis Toolkit (KAT) v2.4.2 [28] and BUSCO v.5.8.3 with the odb12 primates dataset [29].

## 3 Results

The accuracy of our protocol was successfully confirmed across a wide range of reference standards (Figures 3, and 4), demonstrating consistent staining of reference genomes of all sizes.

The genome size of *C. rubicundus* was determined using two standards that were closest in size to the expected primate C-value (inferred from the human genome): *L. niger* and *P. americana* (see Fig. 6).

**Fig. 6.**
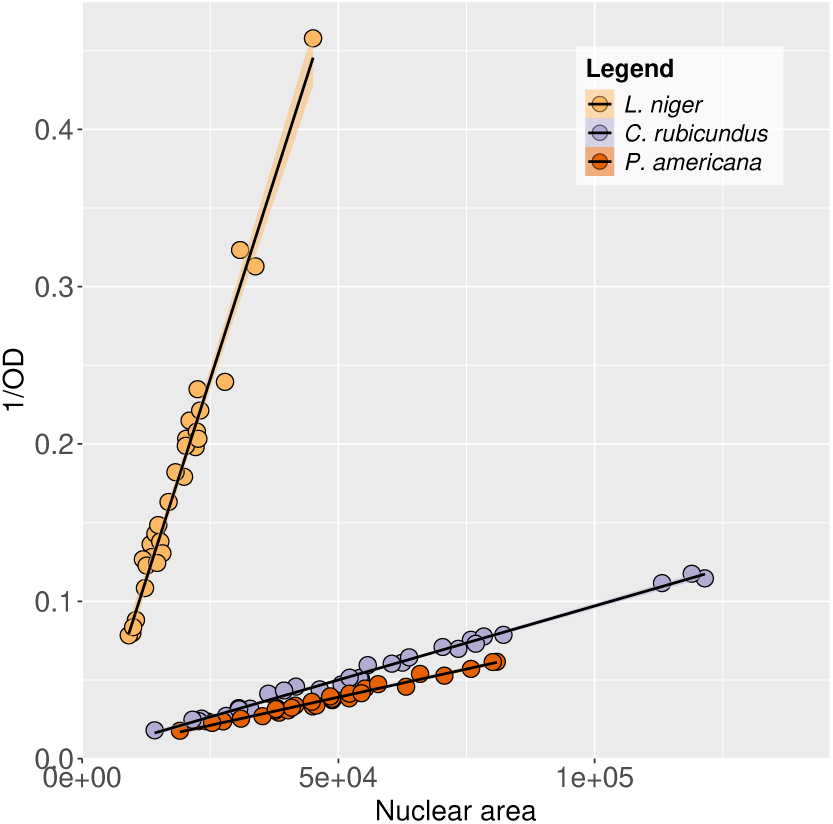
Quality control confirming that the amount of coloration in the G1 nuclei of each species analysed is constant regardless of their area on the microscopy slides. The shaded bands around the regression lines indicate the 95% confidence intervals.

Our Feulgen protocol yielded for each of our three species (our *C. rubicundus* sample as well as our *L. niger* and *P. americana* standards) a large number of nuclei suitable for downstream analyses. Only a few G2 nuclei were detected and discarded for *C. rubicundus* and *P. americana*, whereas they were far more numerous for *L. niger* (presumably before we used near-complete bodies for the ant vs. muscle and brain tissues for the primate and cockroach, respectively). The resulting Feulgen data passed the quality control (Fig 6) and yielded a C-value for *C. rubicunduns* of 2.67 pg (Fig. 7), corresponding to 2.61 Gb (with a CV of 8%). By comparison, k-mer analysis of the Nanopore reads using KAT yielded a genome size estimate of 2.66 Gb, whereas the size of the *de novo* assembly of the same sequence data using Flye was 2.69 Gb. Details of the genome assembly in terms of N50, BUSCO score, and k-mer completeness are presented in Table 1 and Fig. 8, respectively.

**Fig. 7.**
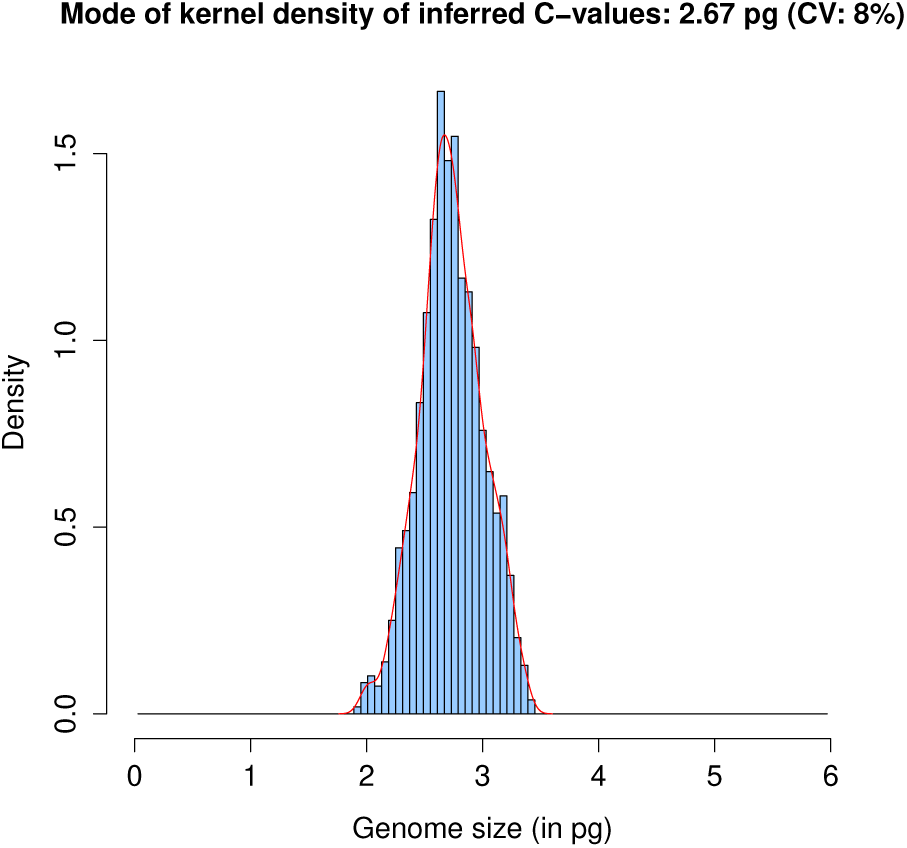
Genome size histogram of *Cacajao rubicundus* with the corresponding Gaussian kernel density distribution (red curve).

**Fig. 8.**
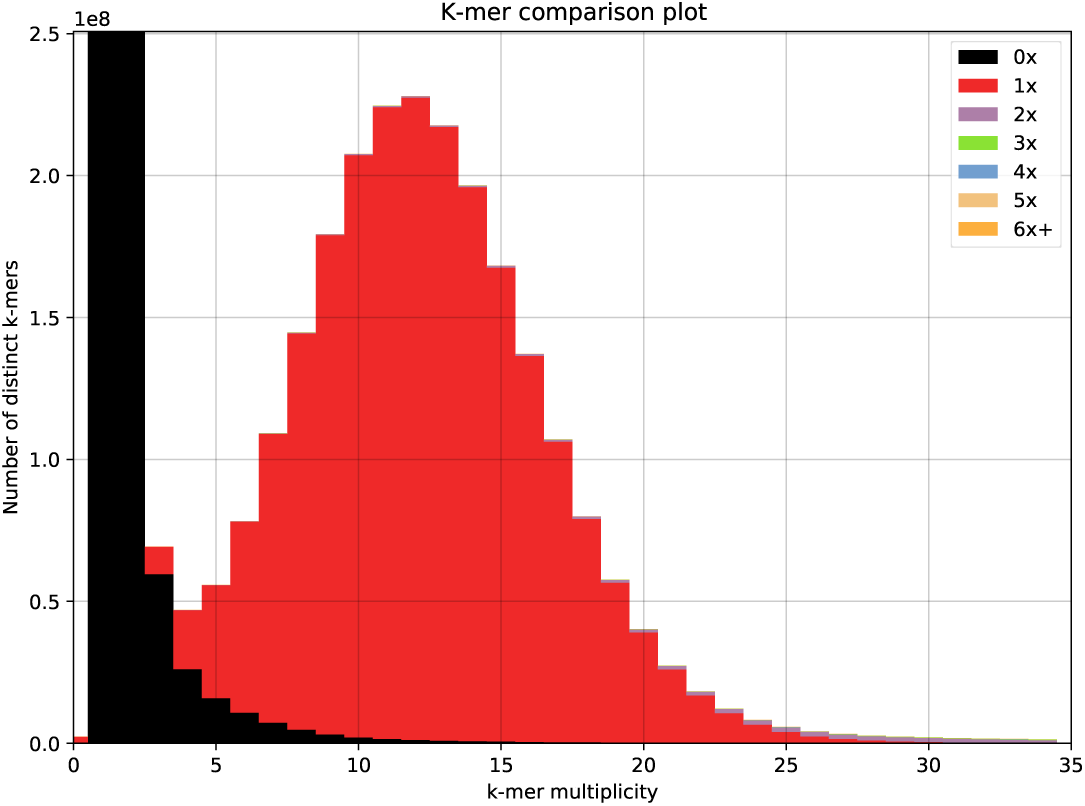
KAT plot of *Cacajao rubicundus* comparing the k-mers in the Nanopore raw reads and in the assembly generated using Flye. Based on the k-mer histogram, KAT estimated the genome size is 2.66 Gb. The legend colors represent the copy number of each k-mer in the final assembly: black indicates k-mers absent from the assembly (0x), red marks k-mers present in a single copy (1x), purple represents k-mers present in two copies (2x), k-mers present in three copies (3x), while all remaining k-mers correspond to highly repetitive copies.

**Table 1.** Assembly statistics for the *C. rubicundus* genome.

| Measure | All scaffolds |
| --- | --- |
| Flye assembly size (bp) | 2,693,375,772 |
| No. of scaffolds | 3165 |
| N50 (bp) | 11,957,981 |
| GC (%) | 40.6% |
| Heterozygosity | 0.12% |
| BUSCO Complete | 99.5% |
| BUSCO Complete and Single | 98.8% |
| BUSCO Complete and Duplicated | 0.7% |
| BUSCO Fragmented | 0.3% |
| BUSCO Missing | 0.2% |

## 4 Discussion

The genome size of *C. rubicundus* was determined as 2.61 Gb using our refined Feulgen protocol, vs. 2.66 using KAT and 2.69 Gb using genome assembly. These values show a very high level of agreement, with the KAT k-mer-based estimate being just 2% higher than the Feulgen estimate, and the genome assembly size being 3% higher than the Feulgen estimate.

A key advantage of the Feulgen method is that it requires very few nuclei (we recommend 30, but a good estimate can already be obtained using as few as 15 nuclei). This is much less than flow cytometry, which typically requires 10^5^ to 10^6^ nuclei [30], hence the Feulgen method appears particularly suitable when working with small organisms or limited tissue samples. Furthermore, the Feulgen reaction is very suited for samples preserved in ethanol (in the case of fresh tissues, the first step in the protocol is actually to plunge them in ethanol).

Besides, flow cytometry is a technique that relies on sophisticated and expensive equipment not ubiquitously available in all laboratory settings. In contrast, the Feulgen method is extremely cost-effective, requiring only a standard microscope, a digital camera, and open-source software. This accessibility makes robust genome size estimation feasible for a broader range of research projects, particularly in laboratories with limited funding.

Our refined protocol incorporates several methodological improvements that significantly improves accuracy and reproducibility, addressing the consistency issues associated with the classical Feulgen protocol [14]. On the laboratory side, there are many factors that can result in problems with Feulgen staining, for instance if the chemical baths are degraded, or if the temperature was not controlled properly during the experiment. In contrast to a different approach that selects only nuclei of similar shapes and sizes, our approach leverages differences of level of DNA condensation among nuclei to confirm, by using nuclei of differences sizes, that the integrated optical density is constant as expected if Feulgen staining was successful (Fig. 3 and 6). Moreover, our method relies on the use of several standards of known genome size to verify that the integrated optical density is indeed proportional to the actual amount of DNA (Fig. 4). Although two standards are sufficient to provide such check, we used here seven standards ranging from the compact genome of *A. vaga* (0.10 pg) to the very large genome of *Scadoxus multiflorus* (44.2 pg) to demonstrate that our protocol yields consistent results across a 442-fold range of genome sizes.

On the computational side, one important refinement we introduced is to estimate the background independently for each nucleus, thereby accommodating local background heterogeneity; by contrast, commercial software to perform Feulgen measurements typically estimate a single global background intensity for the whole slide. Performing localized background subtraction from each nucleus yields more accurate optical density measurements across all the nuclei photographed for a given sample. Furthermore, unlike genome size estimates obtained using the classical Feuglen approach (which are typically reported without standard deviation or coefficient of variation values), our scripts use a Gaussian kernal approximation to calculate the coefficient of variation (CV) for each genome size measurement. Last but not least, our image analysis pipeline takes advantage of open-source tools, specifically R and ImageJ, ensuring complete computational transparency and broad accessibility.

## 5 Conclusion

The refined Feulgen method presented here is a robust, accurate, and economical protocol for estimating genome size. By avoiding the need for advanced equipment, large cell quantities, and exclusively fresh tissue, it represents a powerful and accessible alternative to flow cytometry. Another example of successful application of this method (where it yielded a result consistent both with k-mer spectrum genome size estimates and genome assembly size) was published recently [31]. While our methodology offers a significant cost advantage, its throughput is currently constrained by the time-consuming nature of manual image analysis. We are currently working on automating this process by developing a Python-based pipeline to streamline the entire workflow, from image processing through to the generation of genome size histograms.

## Supporting information

Supplementary File A. Feulgen DNA staining protocol

Supplementary File B. Feulgen image analysis protocol

Supplementary File C. Codes and Excel templates

## Acknowledgments

This study was supported by the Belgian Fonds de la Recherche Scientifique (F.R.S.-FNRS) via “Projet de Recherches” grant T.0078.23 and “Action de Recherche Concertée” to JFF and via “Chargé de Recherches” grant 40017464 to FES. We thank Lucie Vanderlinden for providing the rotifer standard, Clément Thijs and Stéphane Cherrier for providing the insect standards; Marc Reynders and Nanny Mens for providing the plant standards and Olivier De Thier for useful discussions.

---

**Table 1.** Troubleshooting guidelines.

| Step | Problem | Possible reason | Solution |
| --- | --- | --- | --- |
| Microscopy | A pink smear of DNA is observed, without distinct nuclei. | The nuclei were degraded due to poor preservation, or the tissue was excessively chopped. | Use tissues recently preserved in 96% ethanol and keep in freezer till processing; do not chop the tissue too much (at most 5 mins) and avoid using a mortar and pestle. |
| Microscopy | Nuclei are overlapping or aggregated. | The cell suspension was not homogenized well. | Stir the suspension several times using the edge of the same razor blade that was used for chopping. Do this approximately every hour till the drop is completely dry. |
| Microscopy | Crystals and dirt overlap with the nuclei, reducing the quality of the pictures. | The slides were not clean enough. | Before using them, clean the slides with 96% ethanol using a laboratory wipe. |
| Image analysis | In the plot of 1/OD vs nuclear area size, the dots do not form a line passing through the origin. | The staining did not work well, or the nuclei are blurred (not in focus) or there is overlapping coloration from other nuclei. | Use only sharply defined nuclei well in focus and without overlapping coloration for measurements, or use only the largest nuclei (some camera may lack accuracy at measuring the optical density of very dense nuclei). If none of this solves the problem, consider repeating the Feulgen experiment. |
| Data analysis | The R codes fail with an error message complaining about missing columns. | Data were not exported correctly into the tab-delimited txt format expected by the R codes. For example, digits separated by dots in ImageJ become commas when pasted into Excel, which leads to value alteration. | Investigate the exported text files to make sure it is tab-delimited before using it in R. |

**Table 1. Troubleshooting guidelines continued.**
| Step | Problem | Possible reason | Solution |
| --- | --- | --- | --- |
| Genome size calculation | Genome size calculation returns a value several orders of magnitude larger than expected. | Differences in local settings between languages using a decimal point vs. a decimal comma. | Change the Excel settings to decimal dot instead of decimal comma and check the exported text files before running them in R. |
| Genome size calculation | The genome size is much lower than expected. | DNA degradation resulted in poor coloration. | Make sure the tissue samples were preserved in 96° ethanol and kept in a freezer; avoid using samples that are older than 6 months. |
| Genome size calculation | The genome size histogram displays several peaks corresponding to genome sizes that are multiple of each other. | G2 nuclei and/or polyploid nuclei resulting from endoduplication are present in the dataset. | Retain the peak with the smallest genome size, which is most likely the G1 non-polyploid nuclei. If ensure, redo the experiment avoiding tissues likely to contain dividing nuclei (e.g. gut cells) or polyploid nuclei (e.g. heart, and liver in mammals). |
| Genome size calculation | The genome size histogram displays several peaks corresponding to genome sizes that are not multiple of each other. | The inputted genome size of one or more standards is not correct, or their DNA was degraded, or there are nuclei with different genome sizes in the sample. | Check whether the extra peaks disappear when using one standard at time. If so, then one or more standards have a problem - most probably, their expected genome size that was used for calculation is not the actual one, in which case it should be corrected. Alternatively, their DNA might have become degraded: in such case, choose which standard(s) to trust and discard the other(s). If the problem persists even when using only one standard, then the multiple peaks correspond to different populations of nuclei (e.g. host and symbionts), which ought better be distinguished by eye on the slides and analyzed separately. |

**Table 1. Troubleshooting guidelines continued.**
| **Step** | **Problem** | **Possible reason** | **Solution** |
| --- | --- | --- | --- |
| Genome size calculation | Genome size is not accurate (when compared with genome size estimates obtained e.g. from k-mer spectrum analyses). | The standards and samples were not stained or photographed in a consistent way. | Because the Feulgen stain fades through time, standards and samples should be made in the same reagent bath and photographed at the same time. |

**Table 2.**
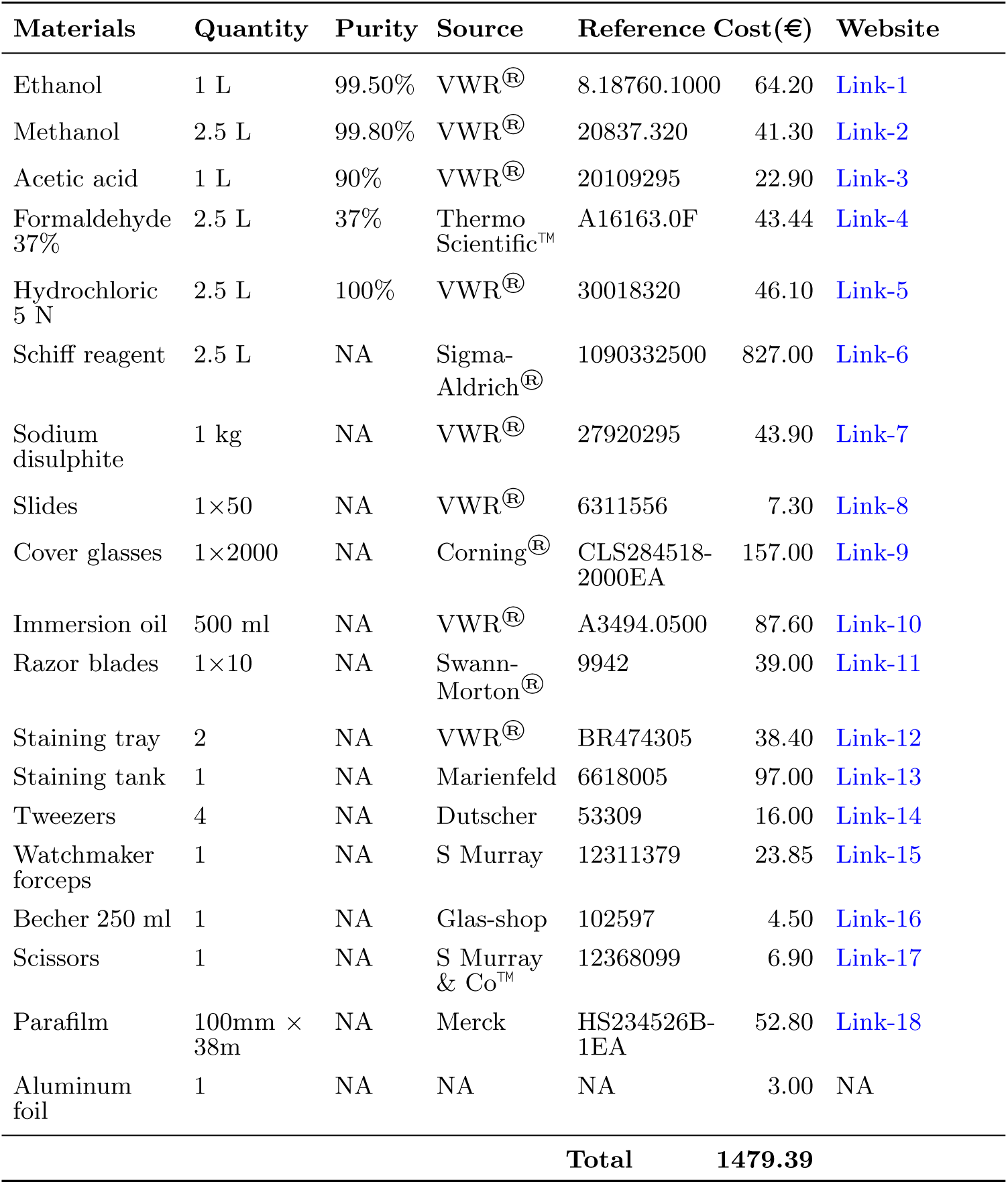
Estimated costs for key reagents and resources of this research.

## Appendix A R codes

R script for generating Figure 3

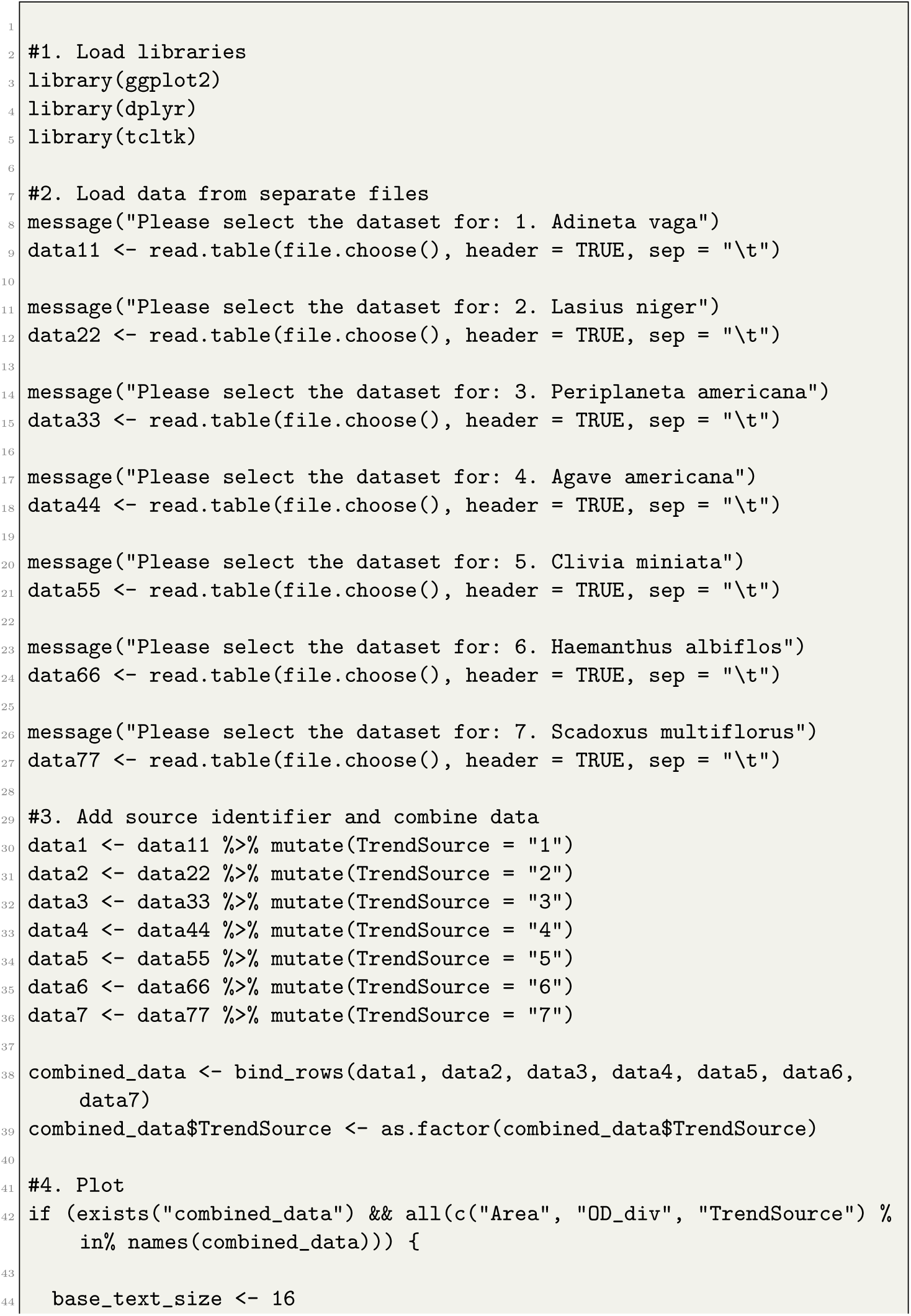
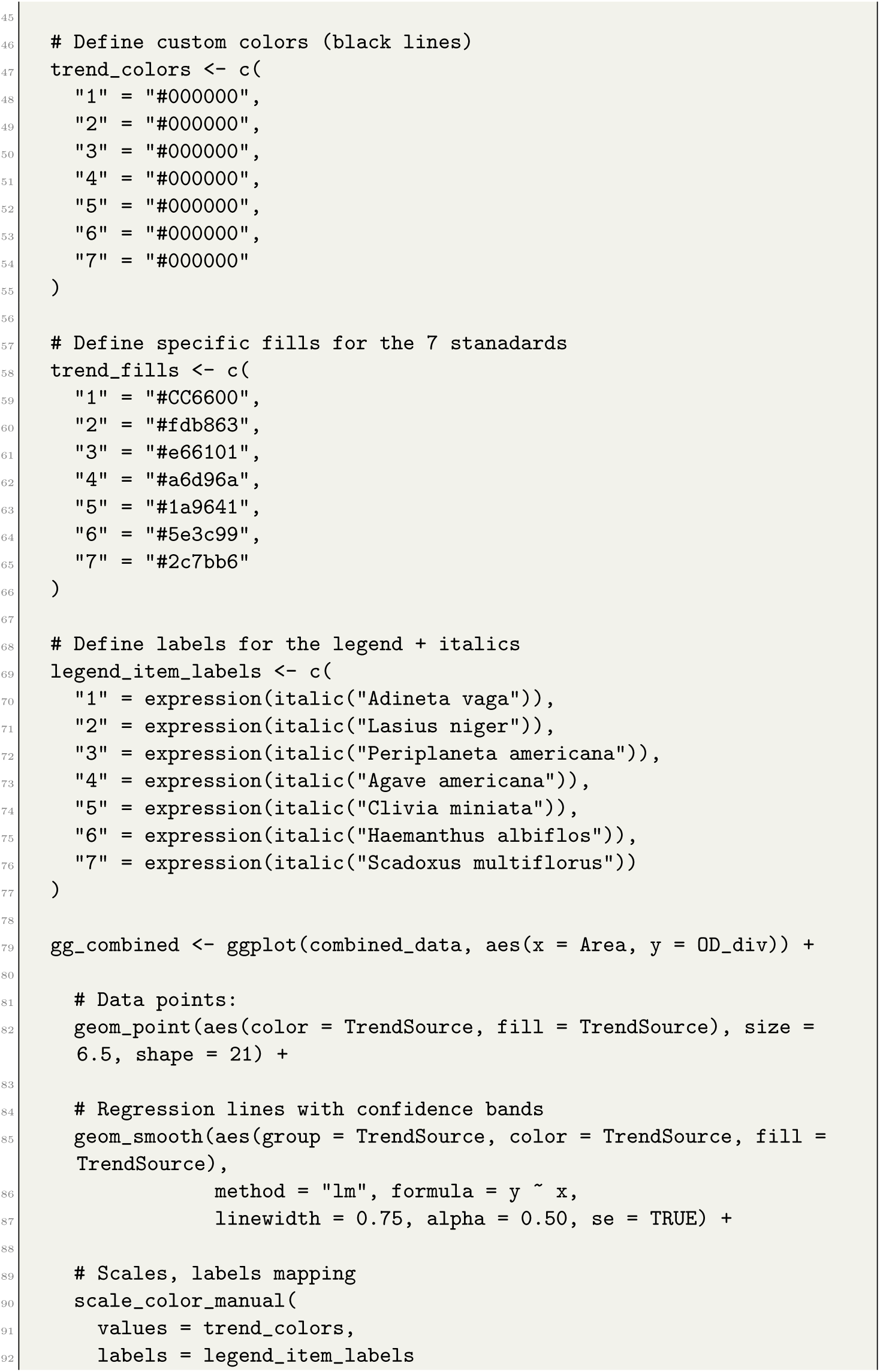
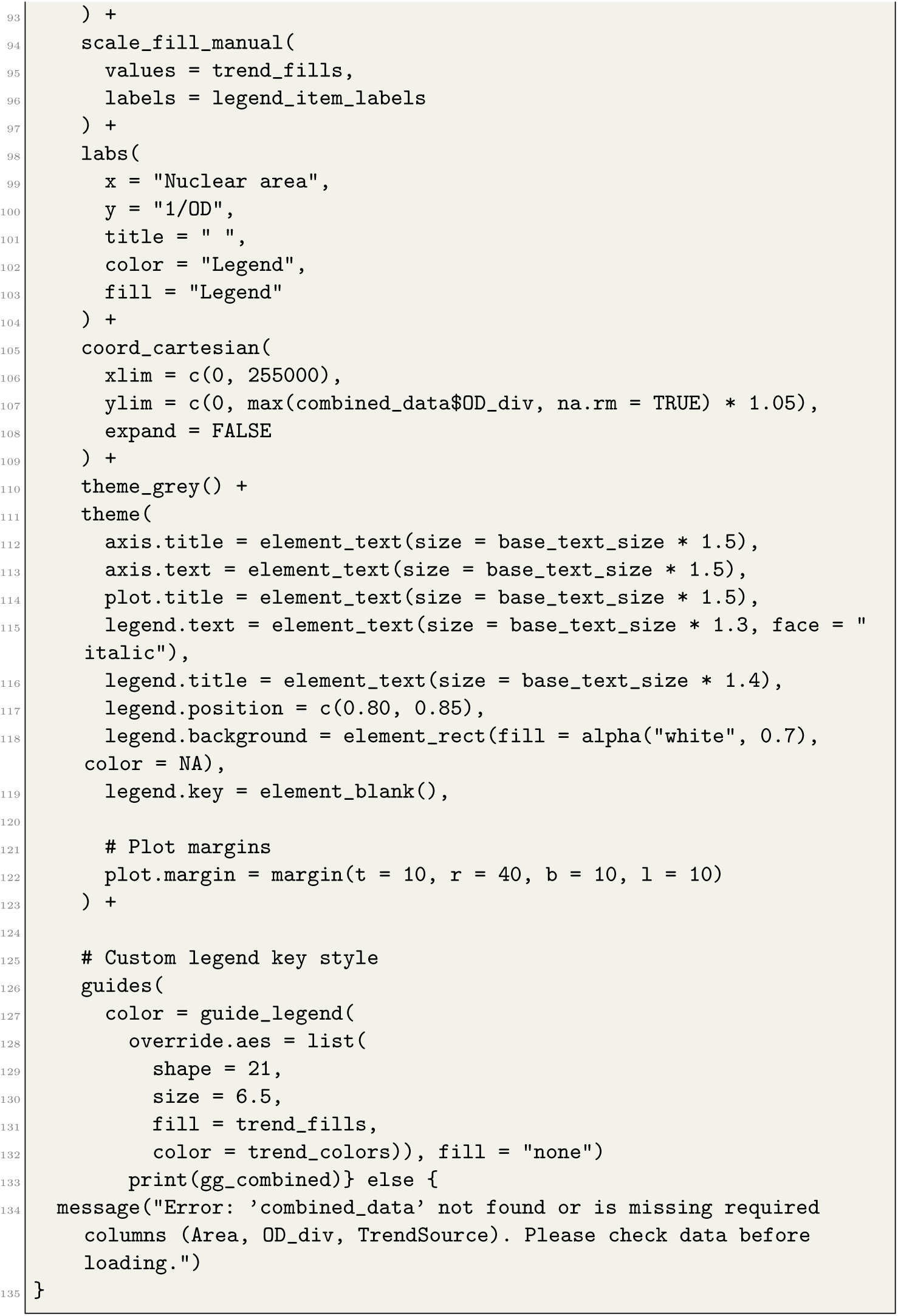

R script for generating Figure 6

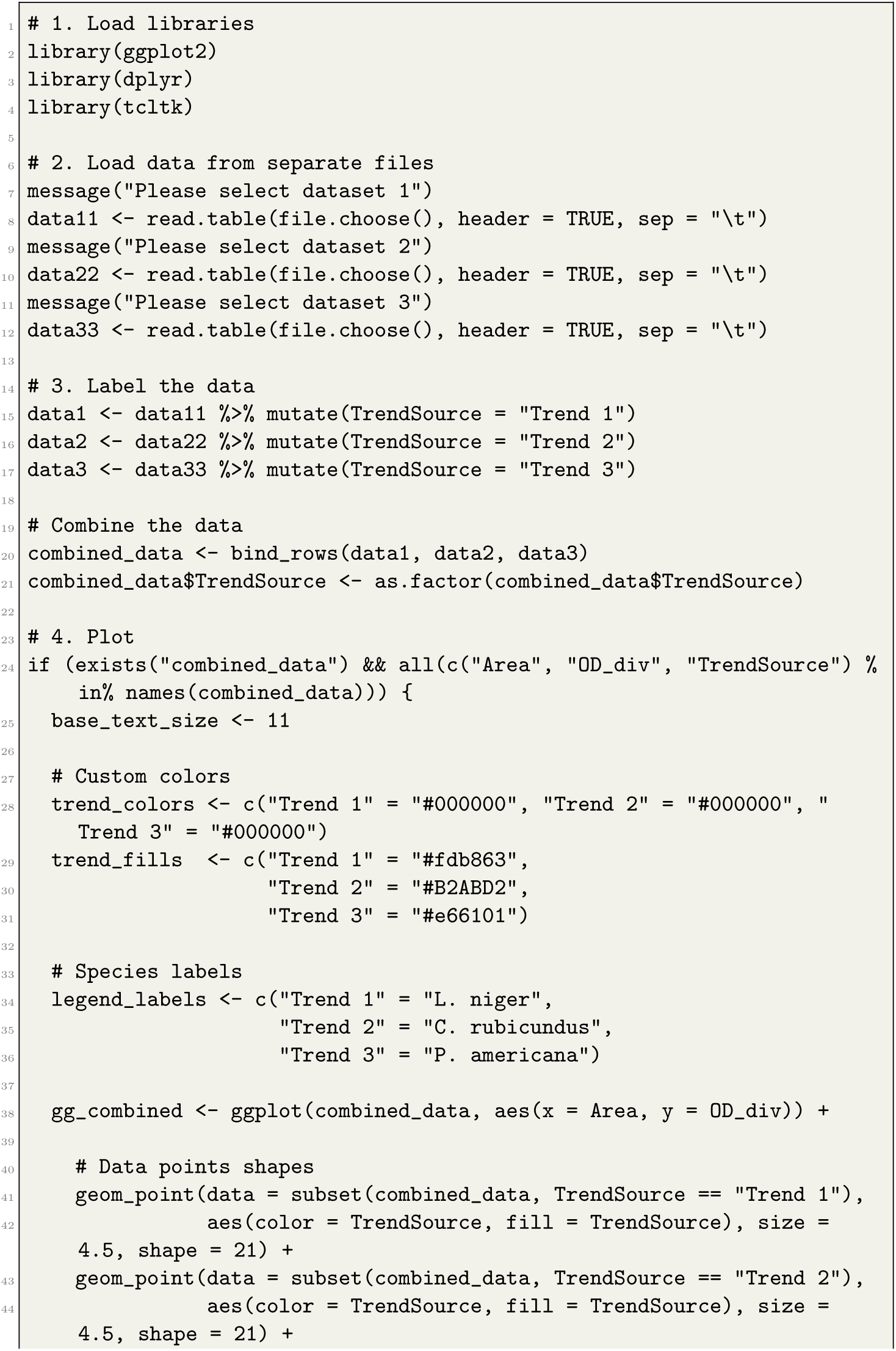
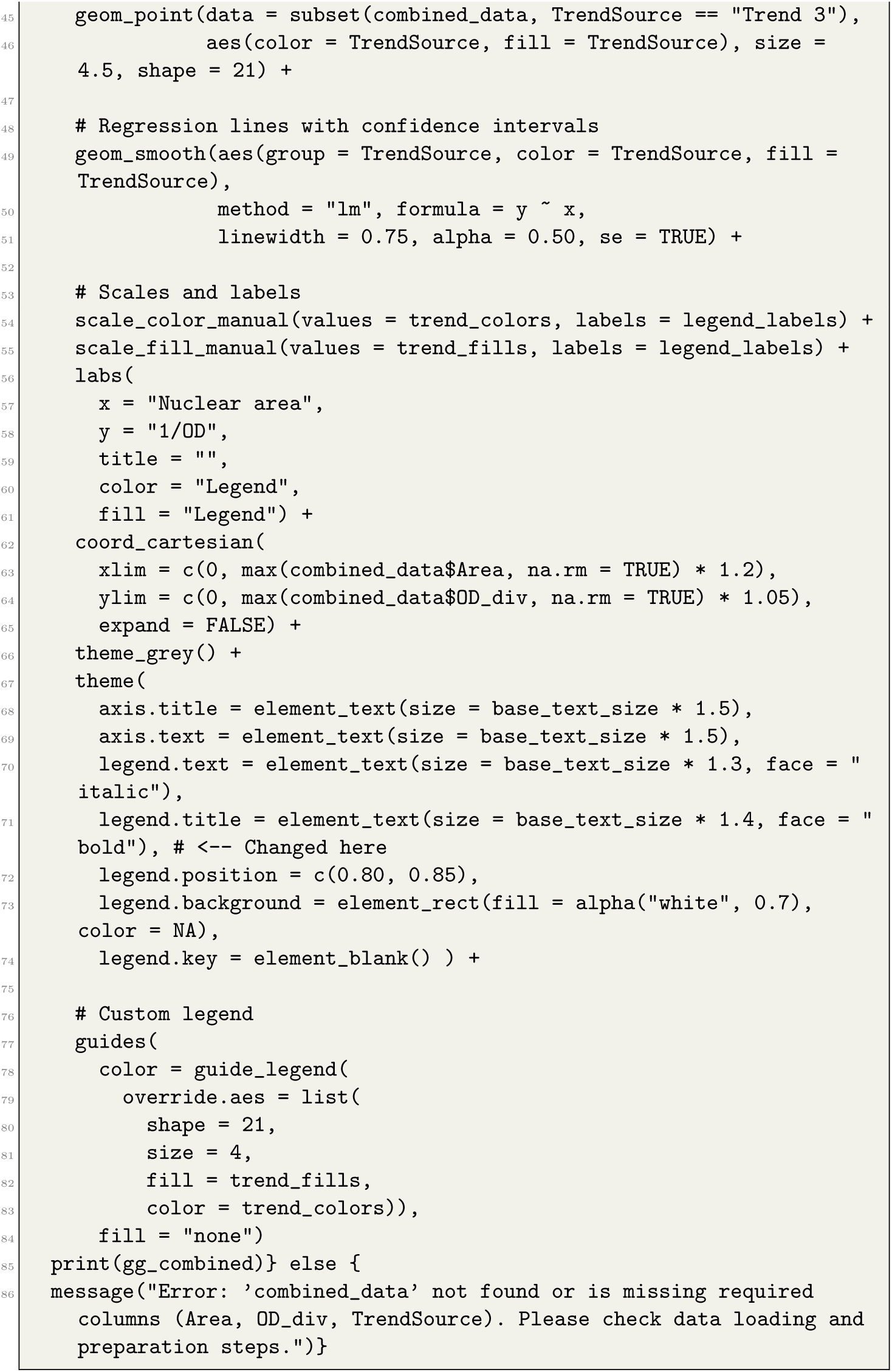

R code for generating Figure 4

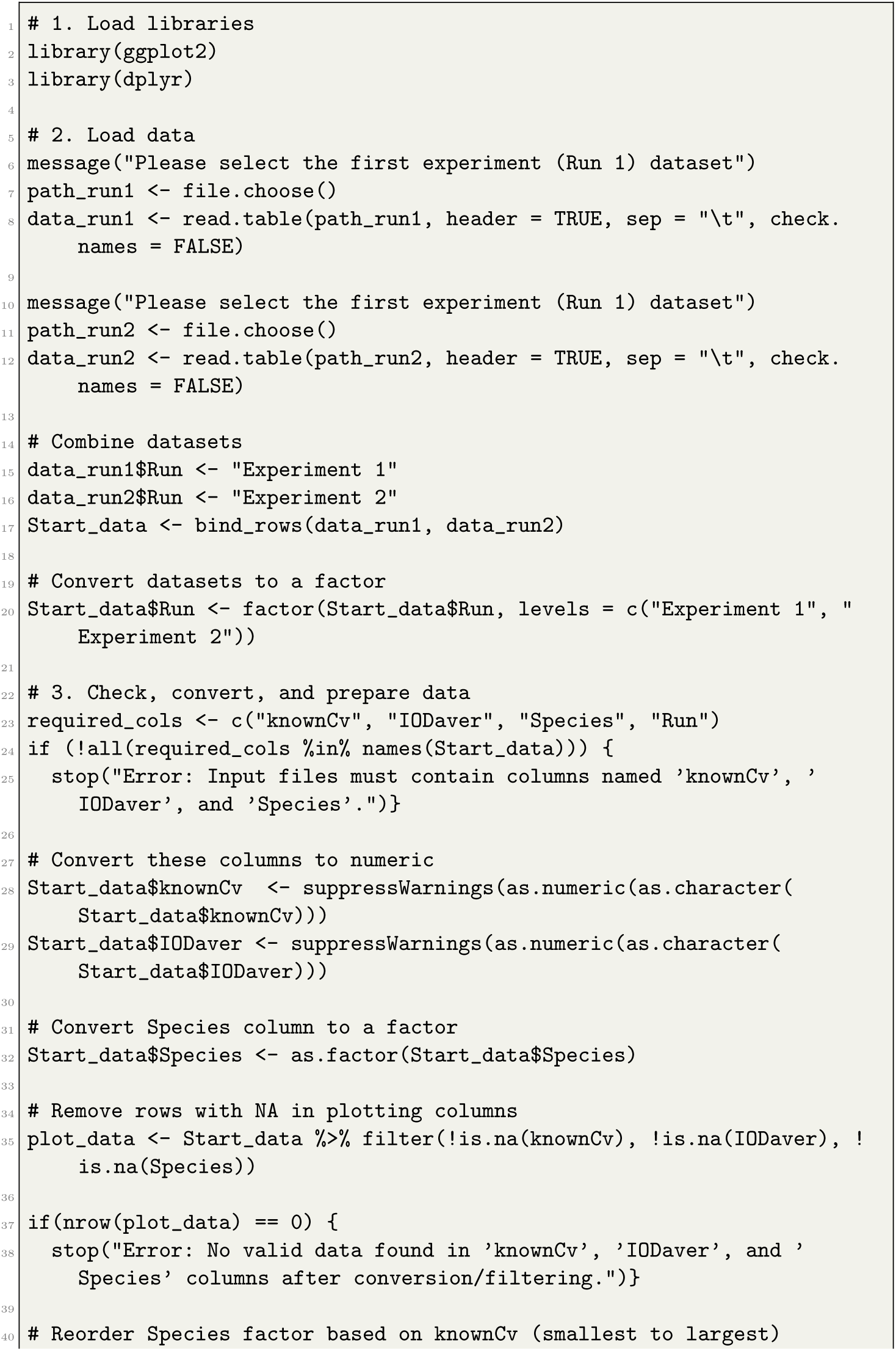
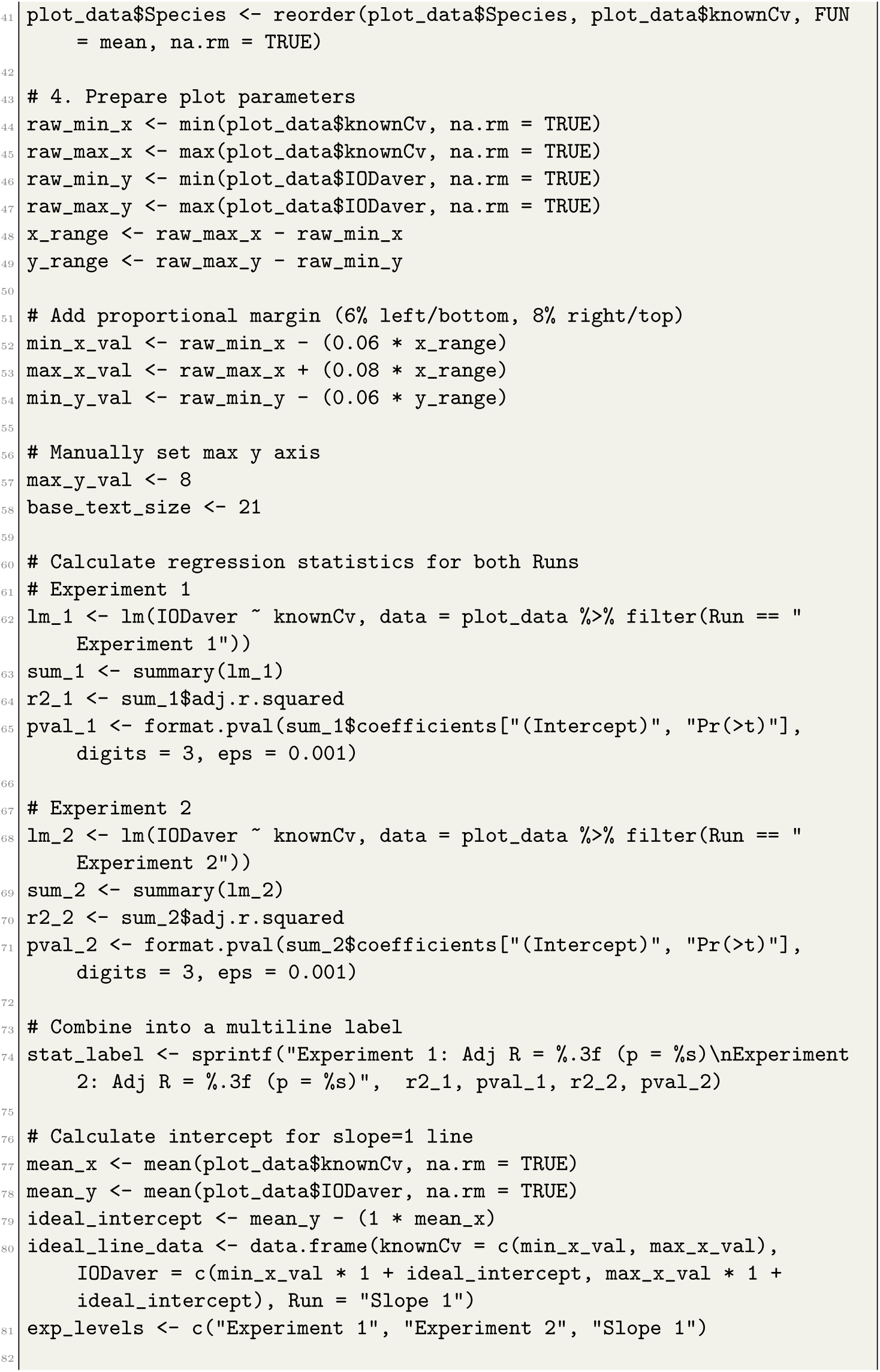
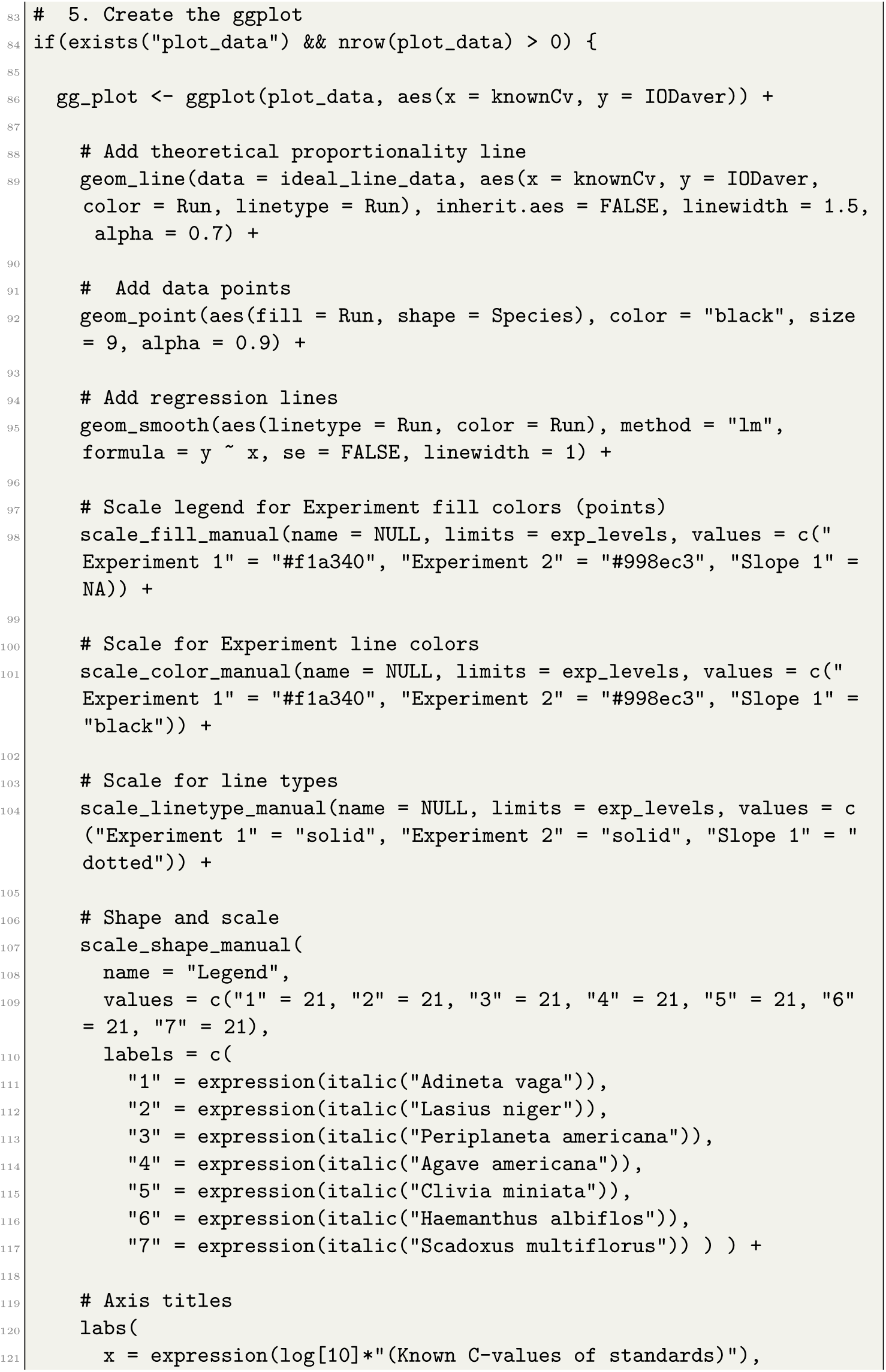
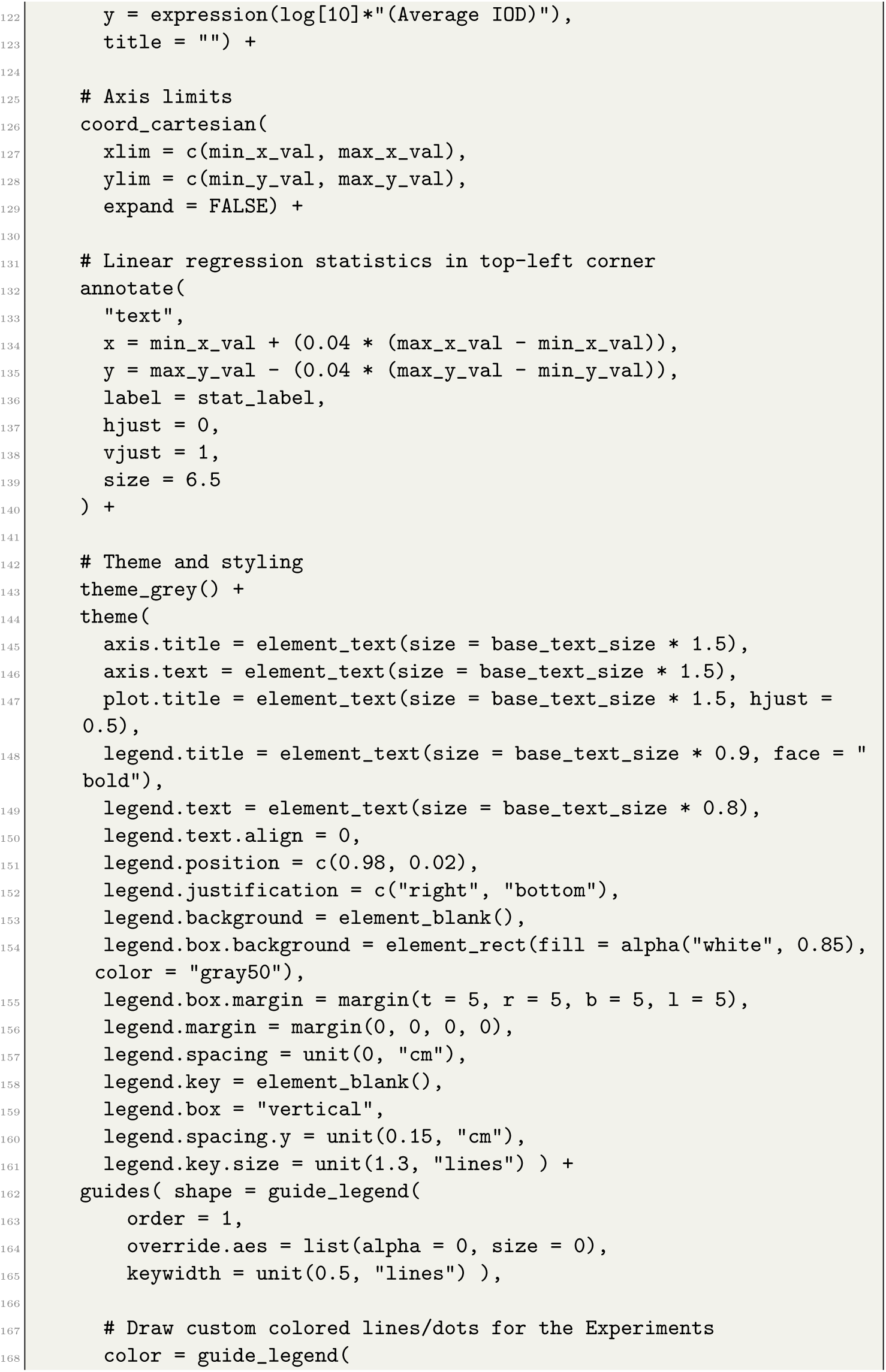
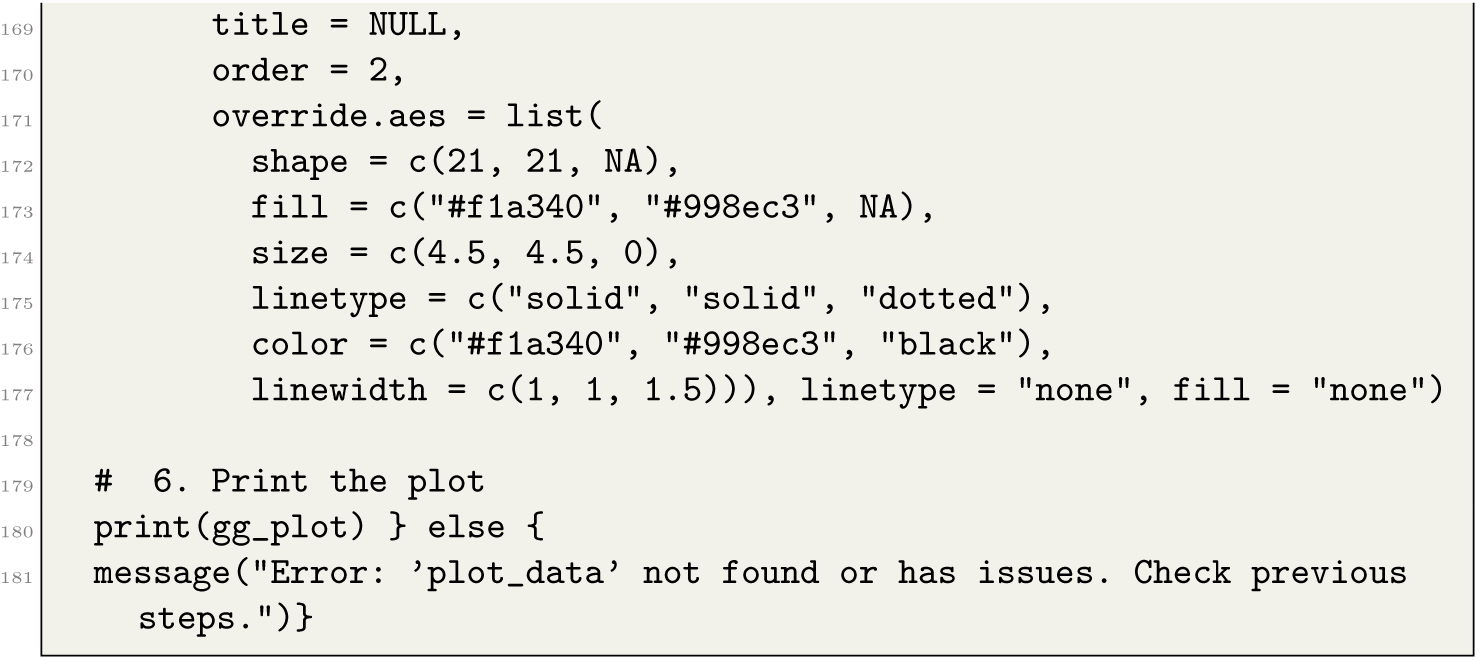

R script for generating Figure 7 (genome size histogram)

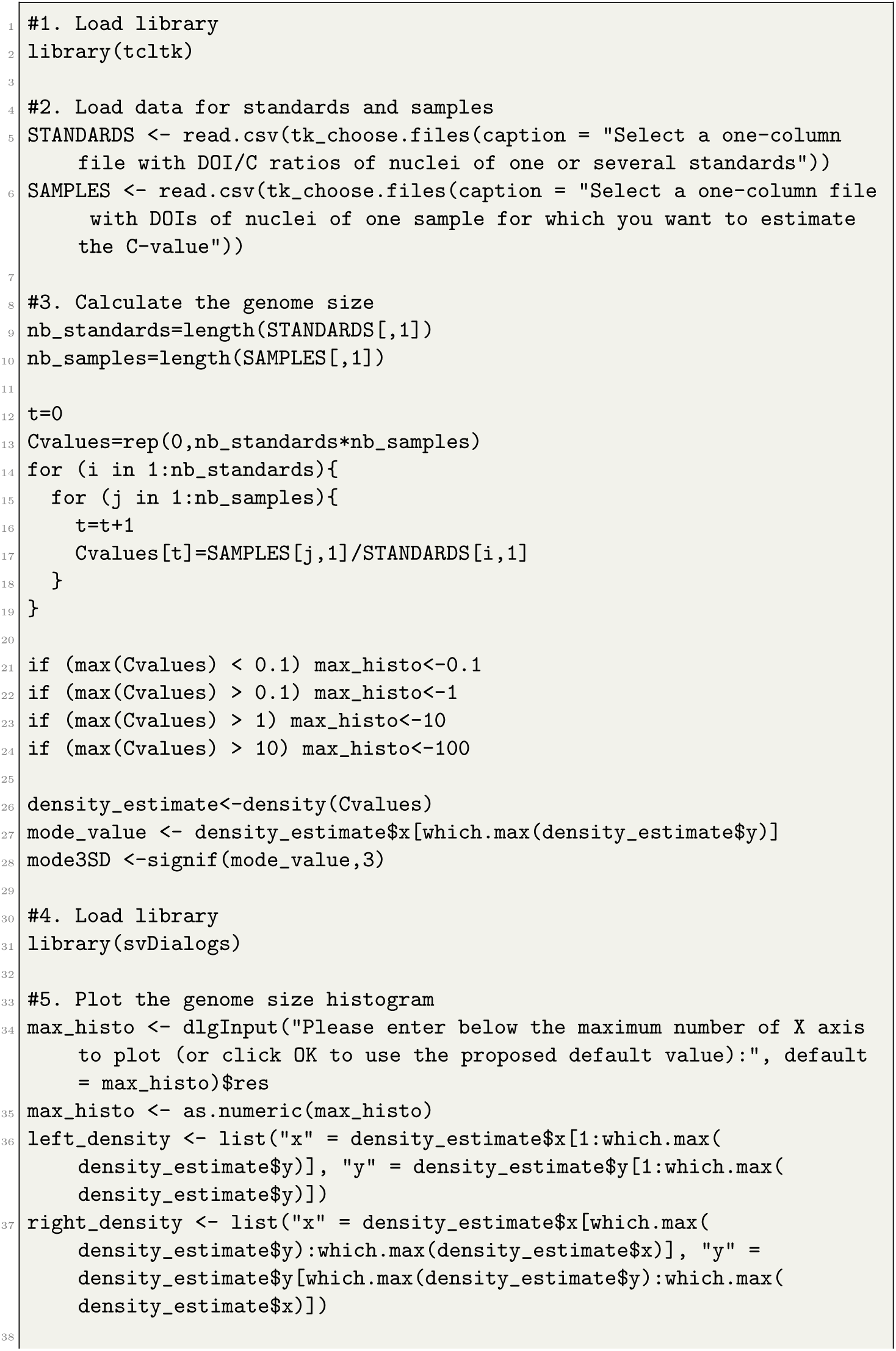
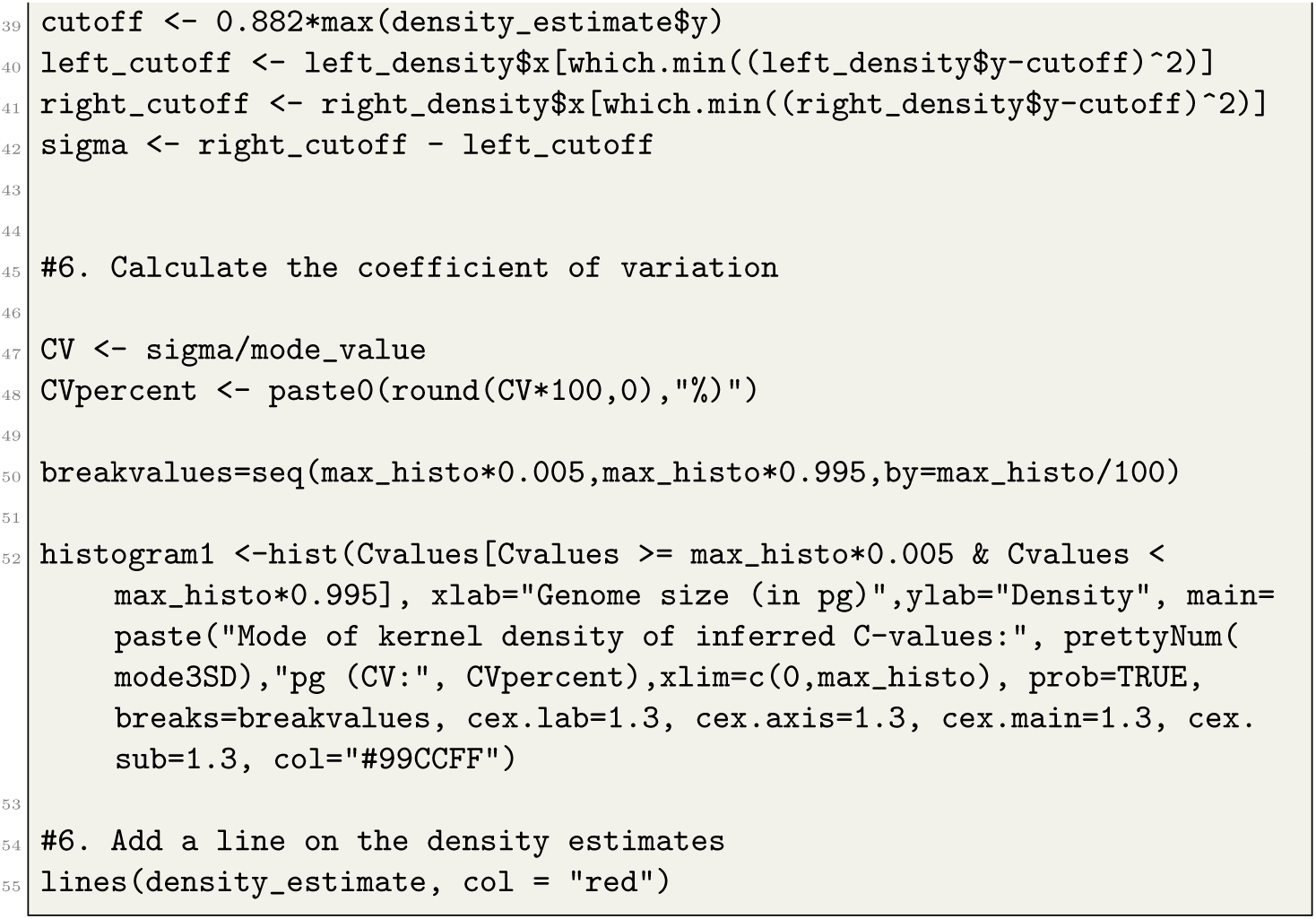

