## Supplementary File A. Feulgen DNA staining protocol for "Refining Feulgen: low-cost and accurate genome size measurements for everyone"

Refined Feulgen Protocol – M.Tawfeeq et al., 2026

Supplementary A – DNA staining checklist

| **Before start checklist** |
|---|
| **Materials** |
| **Biological materials:** |
| ☐ Standards: brain of *Periplaneta americana*; 3 individuals of *Lasius niger* (without abdomen part)<br>☐ Tissues must be fresh or conserved in 96% ethanol at -20°C. |
| **Consumables:** |
| ☐ Ethanol 96% $H_3CCH_2OH$ |
| ☐ Acetic acid 40% $C_2H_4O_2$ |
| ☐ Methanol 99% $H_3COH$ |
| ☐ Formaldehyde 37% (w/w) in aqueous solution stabilised with 7-8% methanol |
| ☐ Acetic acid 90% $H_3CCOOH$ |
| ☐ HCl 5N |
| ☐ Distilled water (DW) |
| ☐ Schiff reagent |
| ☐ Sodium disulphite (SD) $Na_2S_2O_5$, synonymously known as sodium metabisulfite (SMBS) |
| ☐ Plastic based slide staining rack (resistant to chemicals), staining tank or jar |
| ☐ Razor blades, forceps, scissors |
| ☐ Slides, cover slides, petri dish, Becker, funnel, graduated cylinder and bottle |
| ☐ Pipettes and pipette tips P100, P1000 |
| ☐ Immersion oil |
| **Equipment and software:** |
| ☐ Incubator, fume hood, digital balance, Full High Definition camera mounted to compound microscope |
| ☐ Computer preinstalled with the following softwares: ToupView, ImageJ, R studio |

**Feulgen procedure checklist**

| **Step 1: Preparation of tissue samples** |
| --- |
| ☐ Using a graphite pencil, label the frosted end of each microscope slide with the corresponding specimen name. |
| **Note:** Slides must be meticulously cleaned to prevent crystalline artifacts from obscuring nuclei during microscopy in later steps. Immediately prior to use, apply 96% ethanol to the slide and wipe it completely dry with a tissue paper. |
| ☐ Using a sterile razor blade, excise a tissue section with an optimal size of 0.4 cm long × 0.3 cm wide (Figure1, sample 1) from a recommended tissue (e.g., muscle, leg, or brain) and place it directly onto a pre-labeled slide (Figure 1). |
| **Critical note:** Adherence to optimal tissue dimensions (0.4 x 0.3 cm) is essential to yield a suspension of isolated, well-dispersed cells (Figure 1, sample 1). Conversely, processing larger tissue volumes (0.5 cm long × 0.5 cm wide), result in suboptimal dissociation, coarse, granular debris and significant cells aggregation (Figure 1, sample 2). |
| **Note:** For *P. americana* preparation, either immobilize the specimen via direct immersion in absolute ethanol or induce a chill coma by incubating the live specimen at -20°C for 5 minutes. Following that cut the head using scissors, remove the antennas from the head capsule, and apply gentle pressure on the head capsule with forceps to extrude the brain directly onto a microscope slide. |
| ☐ Add 2 - 3 drops of acetic acid 40% then start chopping the tissue directly on the slide using a sterilized razor blade (see Figure 3 for reference). |
| ☐ After chopping the tissues into very tiny particles, add drops of 40% acetic acid to create a suspension on the slide. The resulting suspension solution should be homogenous and not very turbid (Figure 1, sample 1) nor too many grainy textures visible to the naked eye (Figure 1, sample 2). If that's happened, it could be solved by pipetting part of the suspension into a tube then add few more drops of acetic acid on the slide to avoid cell clumps. |
| **Note:** For optimal cell dispersion, use the edge of the razor blade to gently agitate the suspension in a circular motion 2-3 times. This action is crucial to prevent the formation of cell clumps and ensure cells are not overlapped (Figure 1, sample 1). |
| ☐ Keep the processed slide in horizontal position at room temperature for 3 hours until the acetic acid suspension from the previous step is completely dry. |
| ☐ Treat each slide with a few drops of 96% ethanol and allow it to air-dry completely. Subsequently, incubate the slides in the dark at room temperature for 24-48 hours. |
| ☐ Under a chemical fume hood prepare the fixative solution fresh by combining 100% methanol, 37% formaldehyde, and 90% acetic acid in a volumetric ratio of 85:10:5, respectively. |
| **Note:** The fixation reagent must be kept at 25°C overnight before starting the experiment. This preparation should be initiated the evening before starting step 2. |
| **Note:** The total volume of this solution can be scaled proportionally to fit the specific size of the glass tank used. |

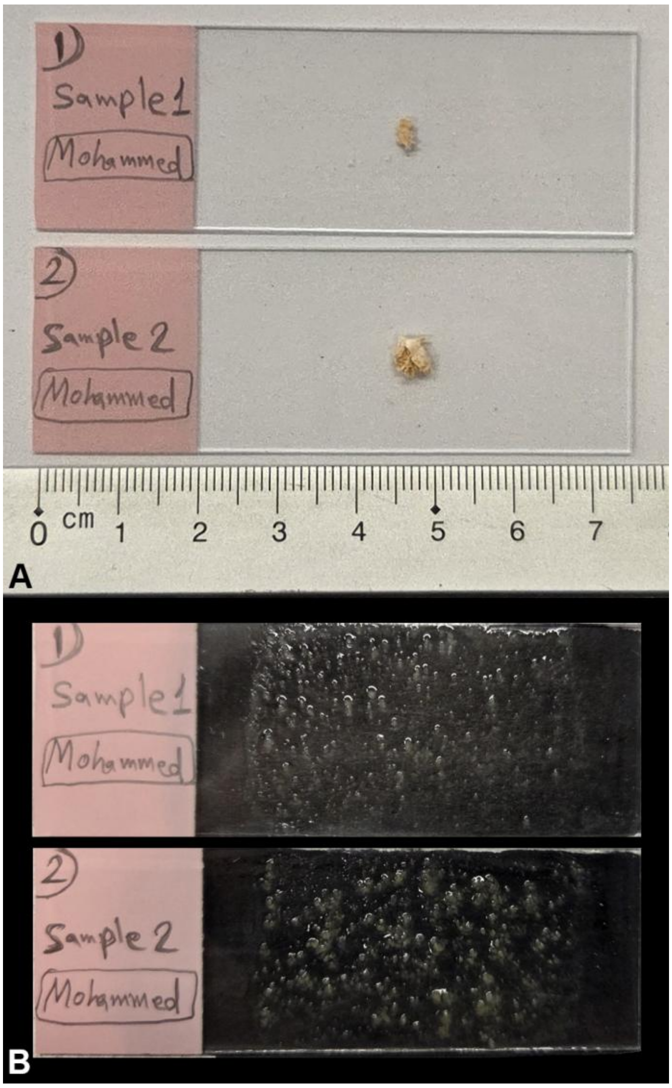


Figure 1. Manual preparation of tissue samples. (A) Dimensions of intact tissue for sample 1 and sample 2, respectively, prior chopping into pieces. (B) Tissue fragments after manual chopping with a sterile razor blade; Sample 1 demonstrates the optimal particle size for Feulgen image analysis.

| **Step 2: Fixation** |
| --- |
| ☐ Transfer the prepared fixative solution into a glass staining tank. |
| ☐ Load the standard and sample slides into a slide rack and immerse it completely in the prepared chemical bath, Figure 2. |
| **Note:** For each Feulgen run, control standards must be processed in the same chemical bath and at the same time as the experimental samples to ensure the validity of the procedure. |
| ☐ Tightly seal the glass tank with its lid, wrap it first with Parafilm and then with aluminum foil to block light, and incubate at 25°C for 24 hours in the dark. |
| **Note:** To ensure efficient chemical reactions, the 5N HCl and Schiff's reagents must be kept at 25°C overnight before starting the experiment. This preparation should be initiated the evening before starting step 3. |

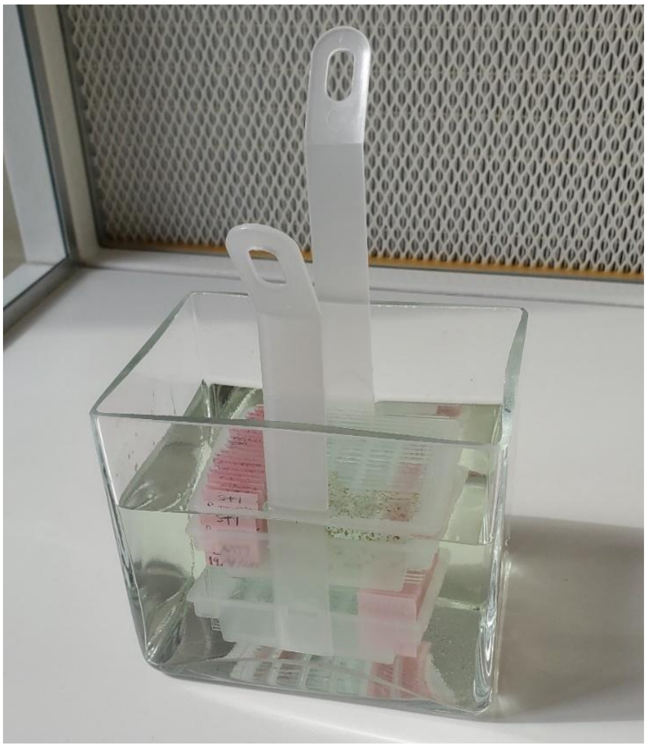

Figure 2. Glass tank contains a slide rack immersed in a chemical bath.

| **Step 3: Hydrolysis and staining** |
| --- |
| **<u>Washing 1 (DW):</u>**<br>☐ Transfer the slide rack into a glass tank containing DW and incubate for 5 minutes at room temperature. Discard the solution and repeat this wash step once using fresh DW. |
| **<u>Hydrolysis:</u>**<br>☐ Place the slide rack in a new glass tank and submerge the slides in 5N HCl. Tightly seal the tank with its lid, wrap with Parafilm and aluminum foil, and incubate in the dark for 2 hours at 25°C. |
| ☐ Repeat washing 1. |
| **<u>Staining:</u>**<br>☐ Immerse the slide rack in Schiff reagent within a glass tank. Seal the tank with its lid, Parafilm, and aluminum foil, then incubate for 2 hours at 25°C in the dark. |
| ☐ During the 2 hours waiting: prepare a fresh solution of sodium disulphite shortly before the 2 hours are over. |
| To prepare 200 ml of solution, mix the following in a glass bottle:<br>☐ Dissolve 1g of SD in 10 ml of distilled water.<br>☐ 188 ml distilled water.<br>☐ 2 ml HCl 5N. |
| **<u>Washing 2 (SD):</u>**<br>☐ Immerse the slide rack in the SD buffer and wash for 5 minutes at room temperature. Discard the buffer and repeat this wash one more time using fresh SD buffer. |
| ☐ Repeat washing 1. |
| ☐ Dehydrate the slides by first applying 70% ethanol for 10 minutes, followed by a final application of 96% ethanol. Allow the slides to air-dry completely. |
| ☐ Incubate the slides in the dark overnight at room temperature. |

| Step 4: Photographing |
|---|
| **Slide preparation for visualization:**<br>☐ Apply a few drops of immersion oil onto the specimen. Gently lower a coverslip at an angle to avoid trapping air bubbles, then add a drop of immersion oil onto the top of the coverslip. |
| **Microscopy:**<br>☐ Place the slide on the microscope stage. Put the 100x oil immersion objective into position and use the fine focus knob to visualize the specimen. |
| **Set illumination:**<br>☐ Set the microscope light intensity to maximum. This illumination setting must be kept consistent for all subsequent image acquisitions. |
| **Set condenser:**<br>☐ Raise the microscope condenser to its uppermost position below the stage to maximize illumination of the specimen. This setting must not be changed during the imaging session. |
| **Camera and software setup:**<br>☐ Connect the camera to the computer via its USB cable and open the ToupView software. |
| **Note:** The camera requires a one-time calibration using a standard slide. This procedure below should be done once only for each new Feulgen run using a corresponding standard slide. |
| **Camera settings and calibration:**<br>☐ After connecting the camera via USB, select it from the "Camera List" window located on the upper left of the software interface (Figure 3) to begin. |

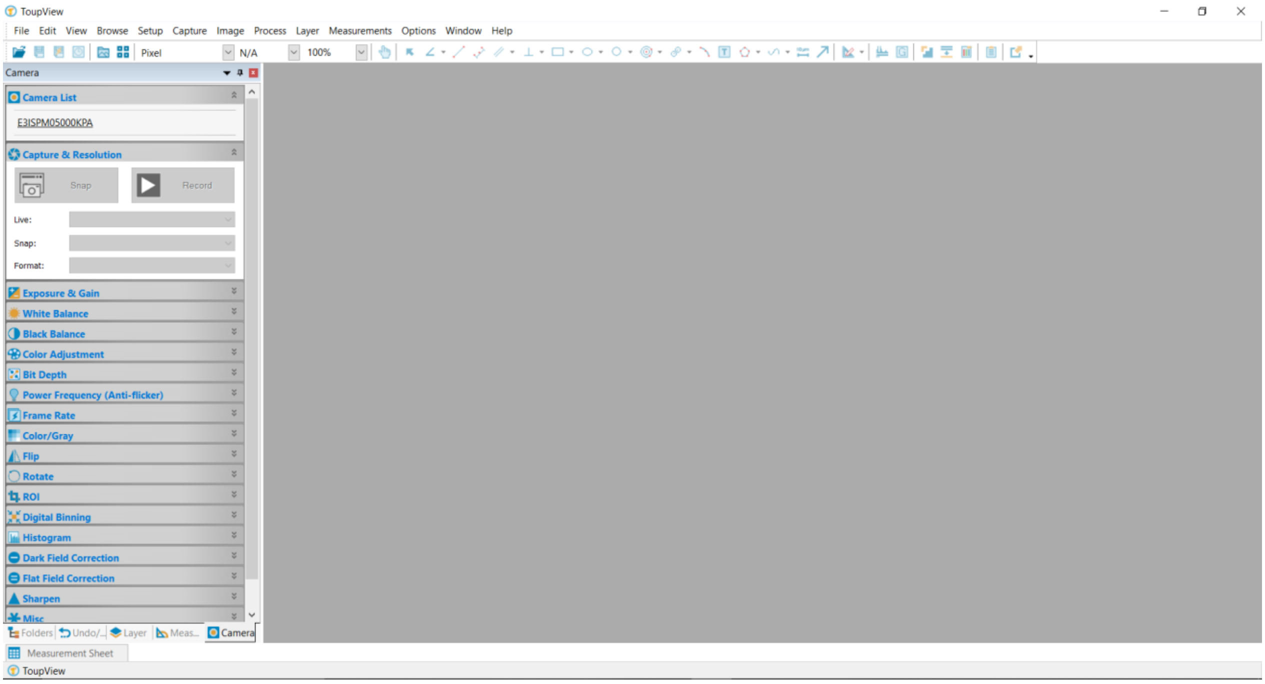


Figure 3

**Initiate live view:**

☐ Click the camera name to display the live video feed from the microscope (Figure 4).

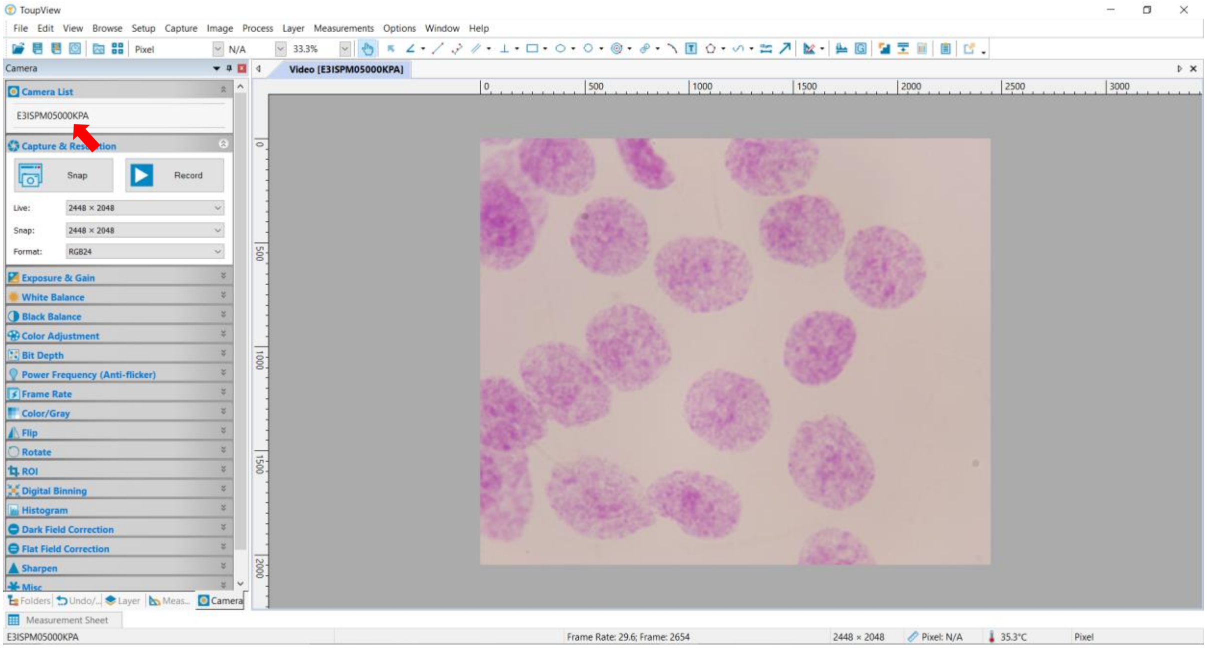


Figure 4

**Set view:**

☐ In the software's zoom settings, select the "Fit to Window" option to correctly scale the live view (see Figure 5).

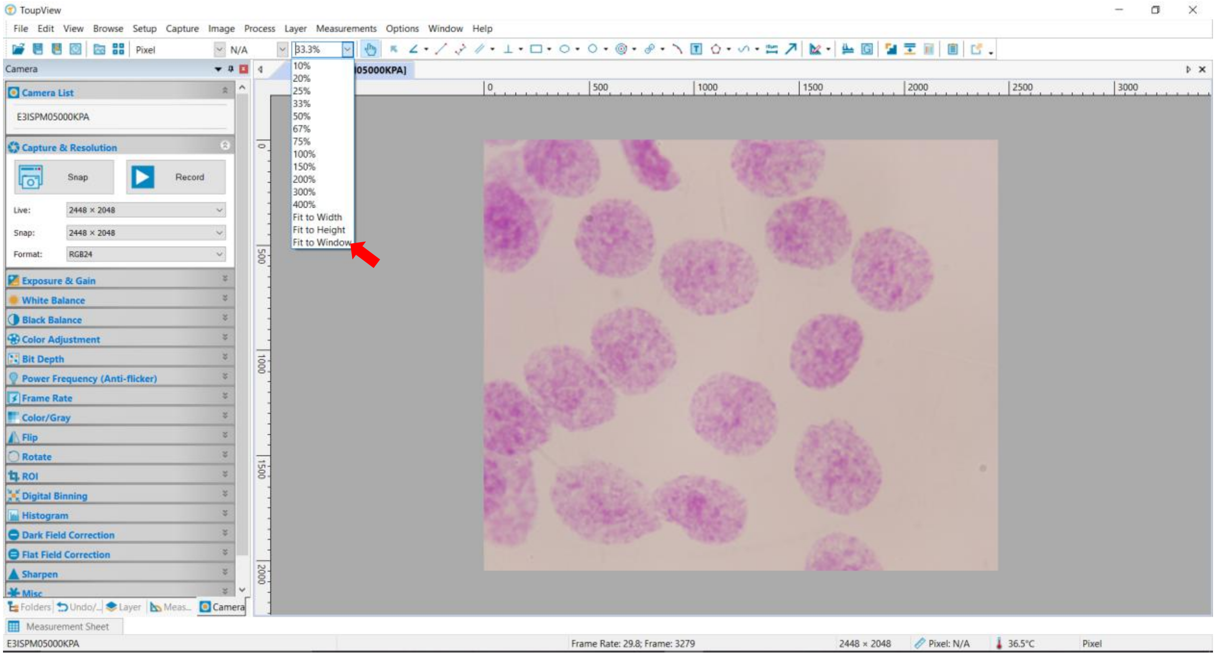


Figure 5

**<u>Set exposure level:</u>**

☐ Click the "Exposure & Gain" button to activate the green selection area. Move this box to a clear region of the slide background, then click "Default" to calibrate the exposure settings (Figure 6).

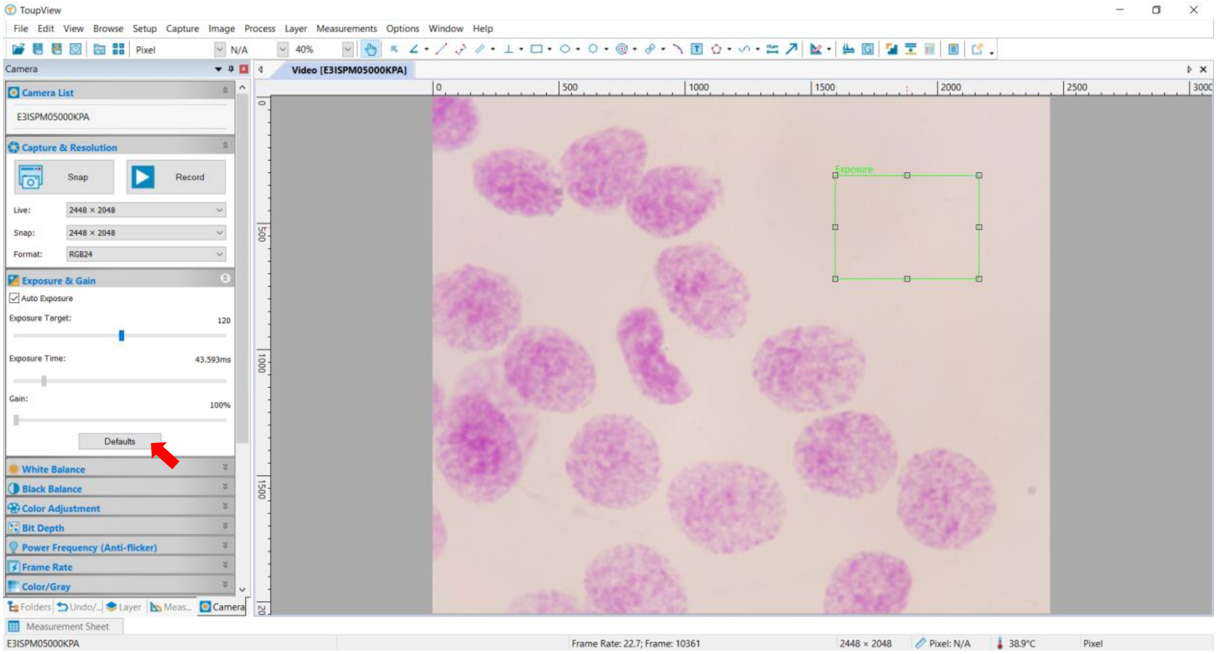


Figure 6

**<u>Lock exposure time:</u>**

☐ Disable the "Auto Exposure" function (Figure 7) to fix the exposure time for all subsequent image captures. Failure to disable auto-exposure will result in variable image brightness and produce erroneous data for genome size analysis.

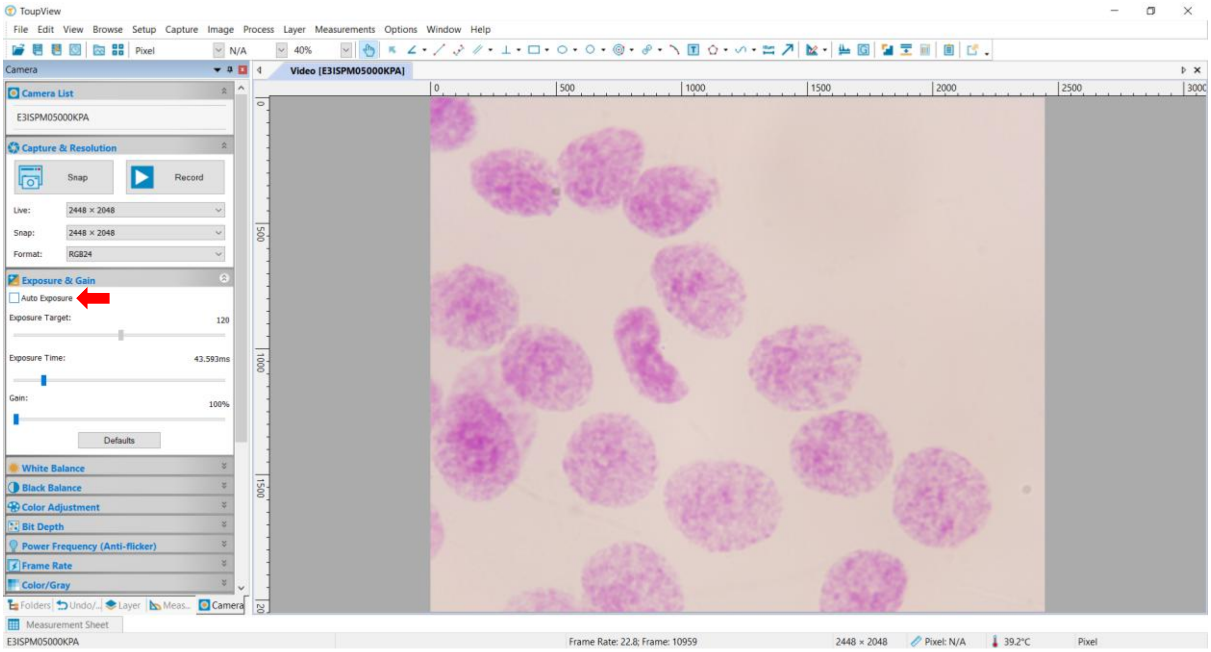


Figure 7

**<u>Set white balance:</u>**

☐ Initiate the white balance calibration by clicking the "White Balance" button. Move the resulting red selection box to a clear background area (Figure 8), then click the "White Balance" button again to apply the setting (Figure 9).

**Note:** To prevent data loss from software errors, manually log the final Exposure Time and White Balance values for each camera calibration per Feulgen experiment. Having these exact settings recorded allows you to seamlessly restore an interrupted session and ensure data consistency

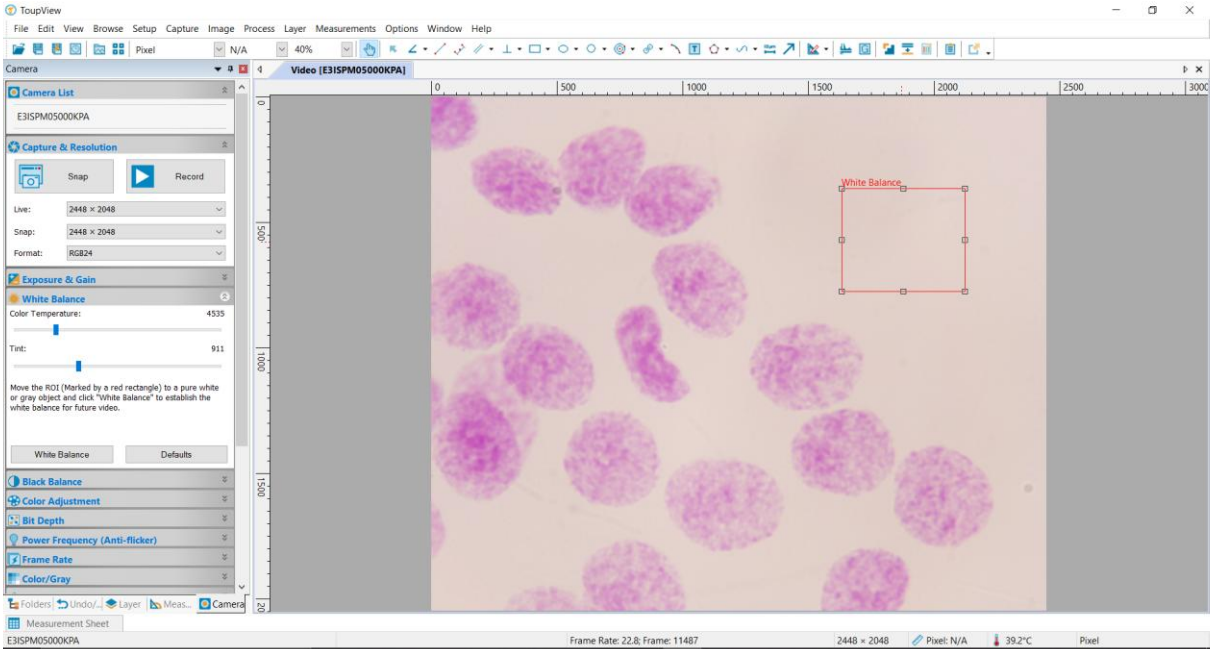


Figure 8

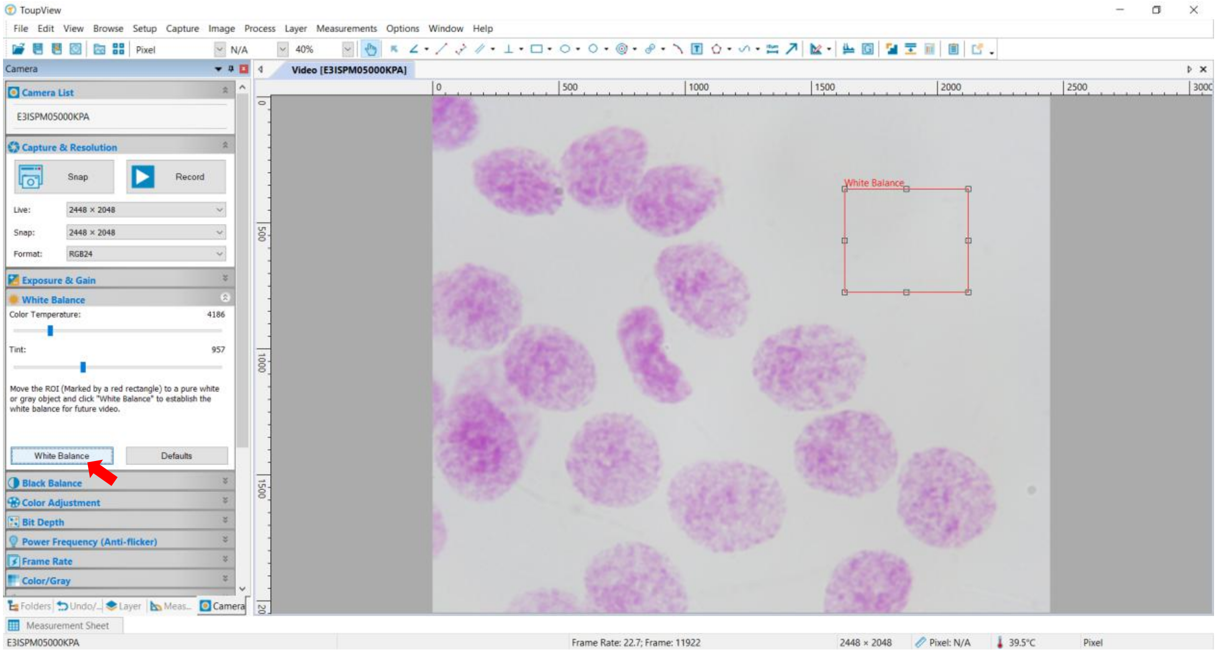


Figure 9

Supplementary A – DNA staining checklist

**Image acquisition:**

☐ Locate a suitable field of nuclei and use the microscope's fine focus knob to achieve a sharp image. Click the "Snap" button to capture an image of the field (Figure 10).

**Critical note:** Standards and samples must be photographed within 2-3 weeks maximum; photographing beyond this period will result in unreliable genome size estimates.

**Image acquisition target:**

☐ Capture 30–50 high-quality images per specimen (see Figure 10) to yield 15–30 measurable intact nuclei required for genome size calculation. Ensure you obtain at least 30 nuclei for the standard reference sample, and a minimum of 15 nuclei for each target sample (aim for 40–80 nuclei for polyploid individuals).

☐ To save all captured images, navigate to File > Save Batch... and choose a destination folder when prompted by the dialog box.

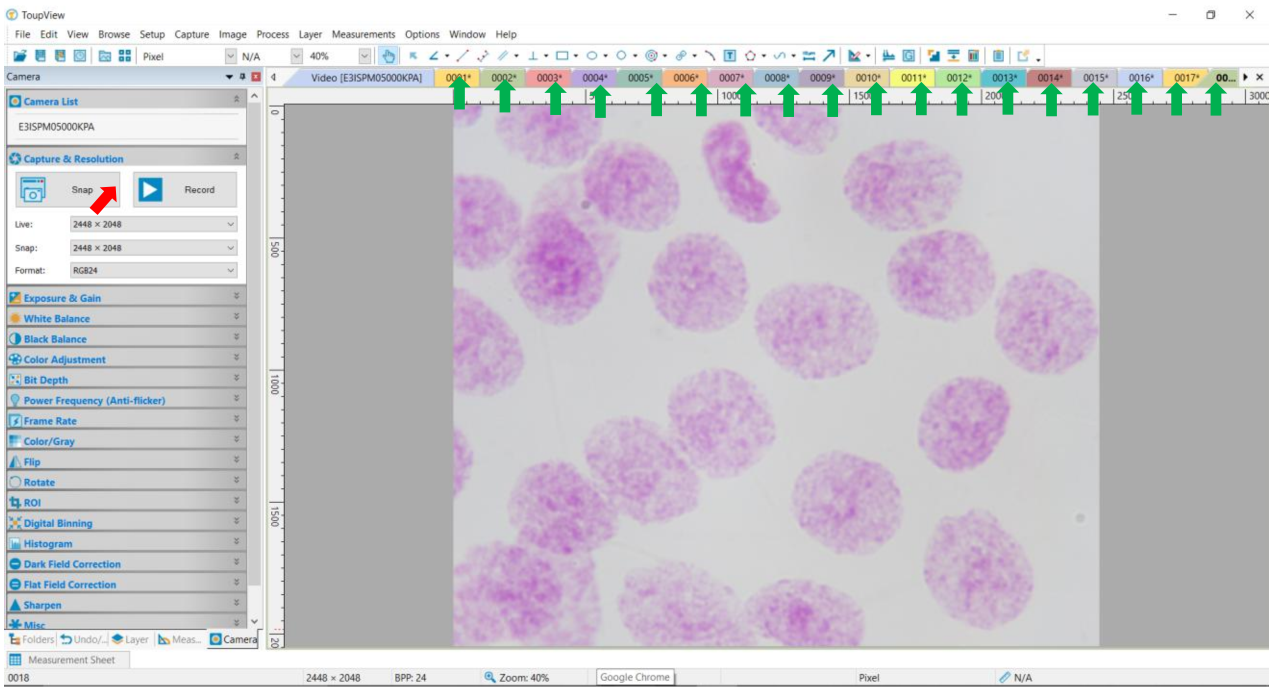


Figure 10

**Critical note:** Before starting the image analysis, quickly check that all the captured images have a homogeneous background luminance intensity. This can be achieved by arranging the images side by side (use e.g., PowerPoint or view large icons in windows system) to make it easier to detect subtle inconsistencies in the background light. If this is the case, it means that the microscope settings were not kept consistent throughout the microscopy session, so it is necessary to re-photograph everything from the beginning.

**Image analysis protocol:**

☐ Go to supplementary B file to follow the protocol of image analysis using the software ImageJ.
