## Supplementary File B. Feulgen image analysis protocol for "Refining Feulgen: low-cost and accurate genome size measurements for everyone"

Supplementary B – Image analysis protocol

| **Step 1: Configure measurement parameters** |
| --- |
| ☐ Launch the ImageJ software. |
| ☐ Navigate to the main menu bar and select Analyze > Set Measurements.... This will open the measurement configuration window (as shown in Figure 1). 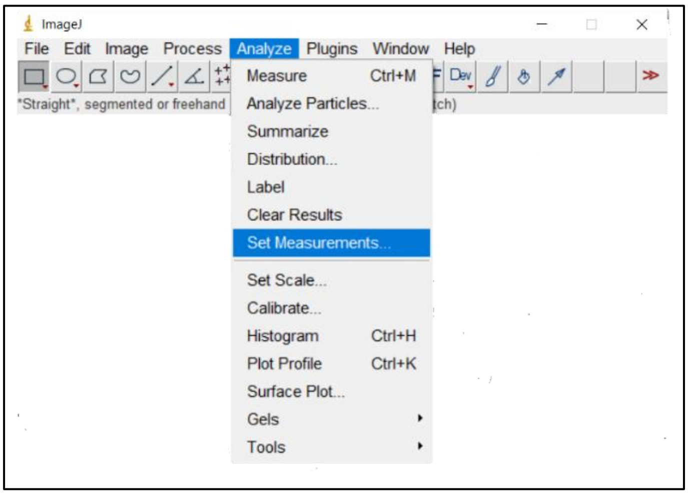 **Figure 1** |
| ☐ In the "Set Measurements" window, enable the "Area" and "Mean gray value" options, then click "OK" to apply the settings (Figure 2). 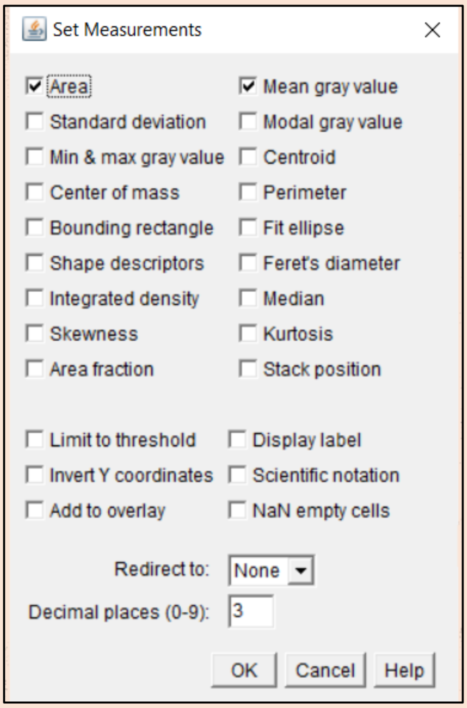 **Figure 2** |

☐ For better visibility during analysis, it is recommended to change the default yellow selection color. This can be modified via Edit > Options > Colors... by selecting a high-contrast color (e.g., blue, red) for the "Selection" tool (Figure 3). Once complete, you may proceed with image analysis.

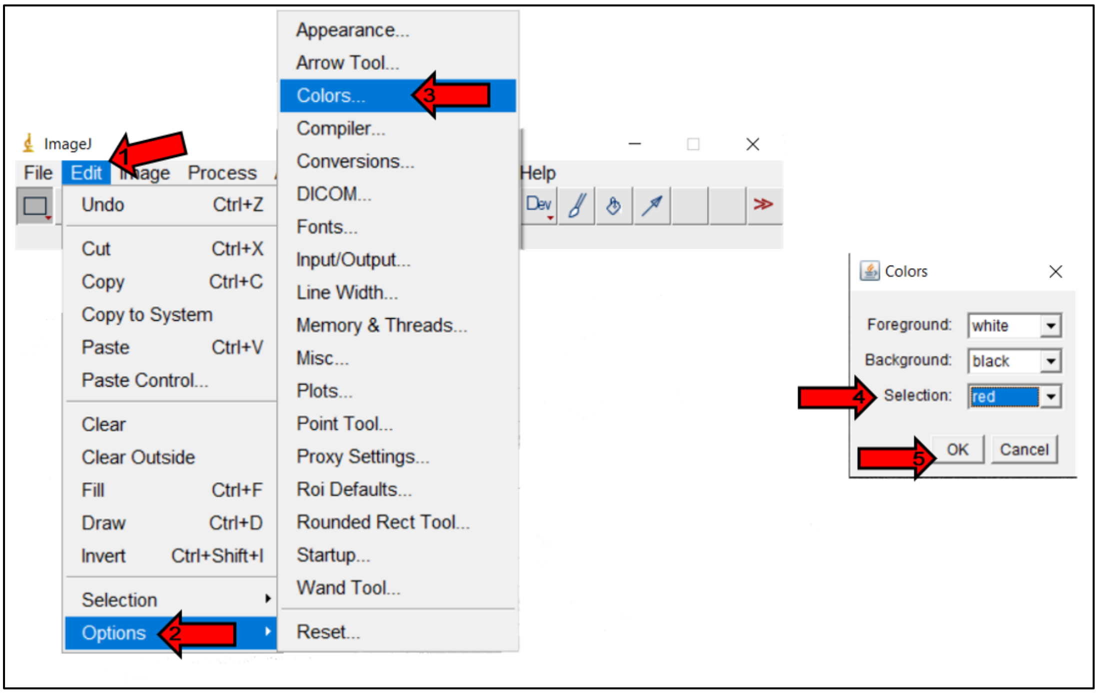


**Figure 3**

**Note:** Firstly, start processing the image set for the standards; then proceed with the analysis of the experimental specimen images.

### Step 2: Data generation

**Loading an image for analysis:**

☐ To open an image for analysis, navigate to the main menu bar and select File > Open.

☐ From the main toolbar, select the "Polygon selections" tool (Figure 4) to begin outlining nuclei.

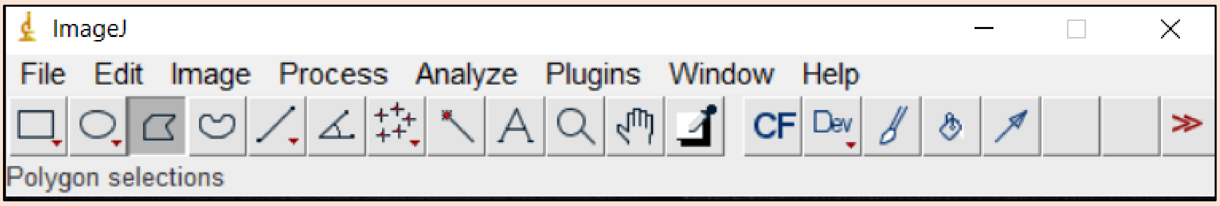


**Figure 4**

**Measuring individual nucleus:**

☐ Zoom and position: hold down the [Ctrl] key and use the mouse scroll wheel to magnify the view, focusing on a single, well-defined nucleus.

☐ Outline the nucleus: with the "Polygon selections" tool active, manually trace the perimeter of the nucleus by placing sequential points with the left mouse button. Continue placing points around the entire boundary until you return to the starting point to form a closed, continuous selection (as depicted in Figure 5).

☐ Record the measurement: with the nucleus selected, navigate to the main menu and select Analyze > Measure or [Ctrl] + [M] (Figure 6). The "Area" and "Mean gray value" for the selected nucleus will automatically populate in the "Results" window (Figure 7).

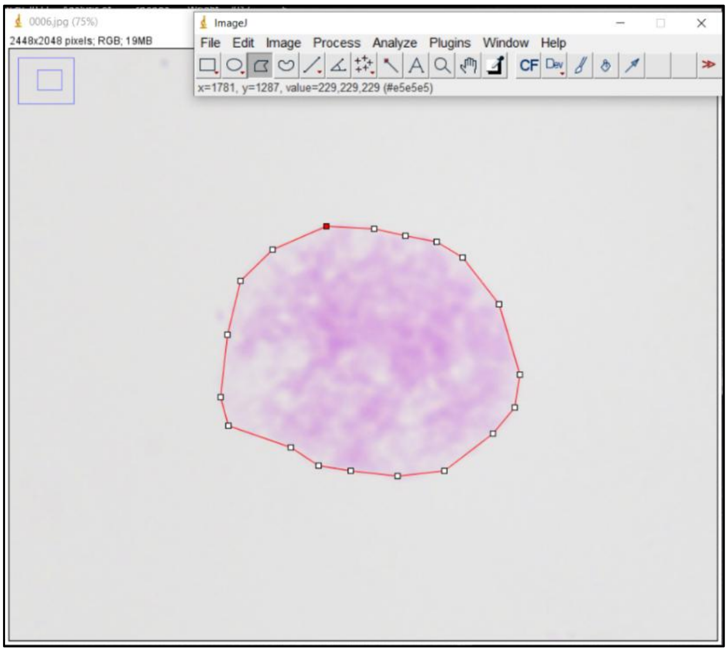


**Figure 5**

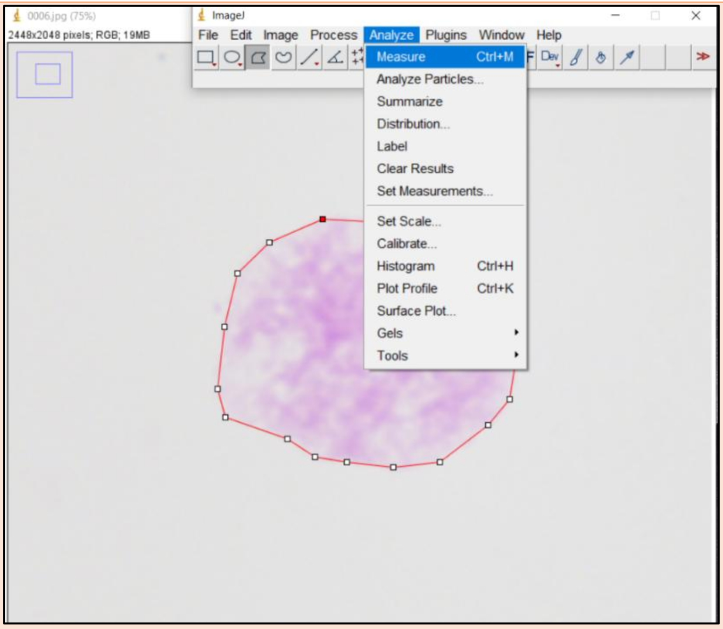


**Figure 6**

**<u>Measure Local Background of the Nucleus:</u>**

☐ For each nucleus measured, use the "Polygon selections" tool to select an adjacent background area to form an enclosed shape similar to donuts shape (Figure 7). Record its parameters by selecting Analyze > Measure or pressing [Ctrl] + [M].

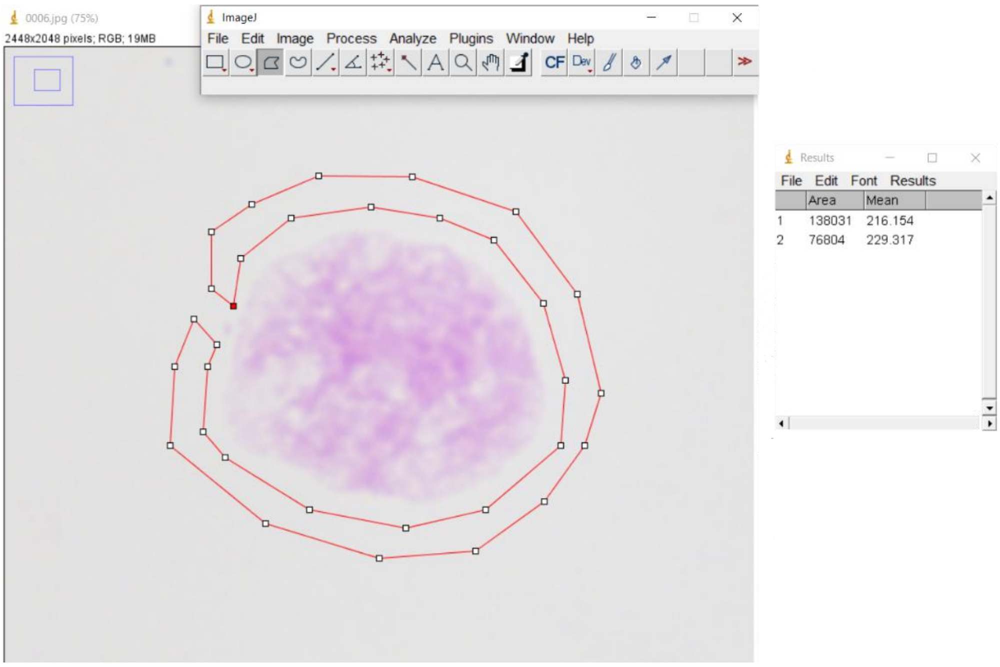


**Figure 7**

**Note:** Measuring a single nucleus generates two rows in the results table (Figure 7). The first row provides the nucleus parameters (Area and Mean Gray Value), while the second row provides the corresponding background data. Only the Mean Gray Value from the second row is used for analysis; disregard the background area measurement.

**Transferring measured data:**

☐ Copy all data from the "Results" window and paste it into the corresponding columns of the "image_analysis_standards_template.xlsx" and "image_analysis_sample_template.xlsx" files, which are provided in the supplementary materials.

**Target nuclei count for analysis:**

☐ Repeat the measurement process until data has been acquired for 30 different nuclei from each standard and for 15 different nuclei from each experimental specimen.

**Notes:**

- Automatic calculations: The provided Excel template automatically computes Optical Density (OD), Integrated Optical Density (IOD), and IODC values from your raw data.
- Standard C-value adjustment: Update the standard C-value in the formula cells to match your specific reference standard before proceeding.
- Quality control check: After entering your measurements (30 nuclei for the standard, 15 per specimen), inspect the template's scatter plot of 1/OD versus nuclear area. A linear relationship is required; if linearity is missing, refer to the troubleshooting guide before continuing further.

### Step 3: Linear regression models

**Preparing data for R:**

**First pre-test**

☐ For each standard and specimen, create a new two-column, tab-delimited text file containing the Nuclear Area and 1/OD data.

☐ Save this new worksheet as a tab-delimited text file (.txt). This file will serve as the input for the R script.

**Executing the R script:**

☐ Using the provided R script ("LM_model.R"), run a linear regression on this data file. The script will generate a statistical summary and a plot. You may need to adjust the text coordinates in R script to ensure they are placed clearly on the plot. The final output should resemble the example shown in Figure 8.

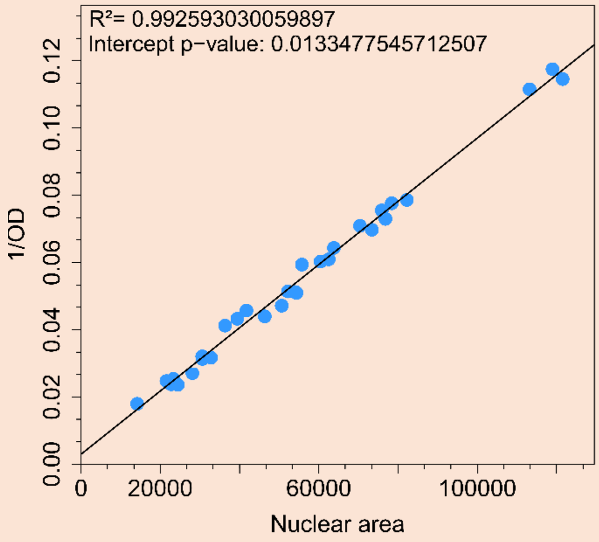


Figure 8

**Note:** The previous described processes in this step must be repeated for every individual standard and every individual specimen.

**Second pre-test**

☐ Populate the " LM_known_C-value_template.xlsx" template with the mean IOD for each standard along with its C-value and save it as a tab-delimited text file.

☐ Using this text file as input, execute the " LM_known_C-value.R" script to generate the linear regression model and calibration plot (see example Figure 9).

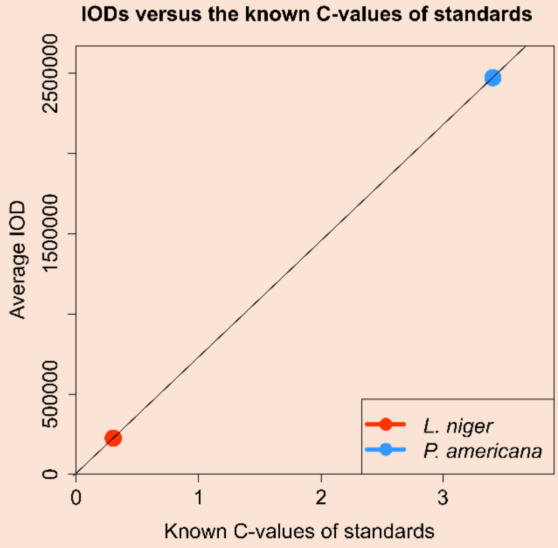


**Figure 9**

**Note:** After verifying the standards data via the two pre-tests, process samples of undetermined genome size by repeating step 2, followed by the first pre-test only described in step 3.

### Step 4: Generating genome size histogram

**Exporting standards' data:**

☐ In a new worksheet, aggregate the IODC values from all standards into a single column (a total of 60 values when using two standards). Export this consolidated data by saving the sheet as a tab-delimited text file.

**Exporting sample's data:**

☐ For each experimental specimen, copy its IODs values into a new worksheet and save it as a tab delimited text as well.

**Generating the genome size histogram:**

☐ Execute the R script named "Genome_size_histogram.R", which is available in the supplementary materials.

The script will prompt you to load data in a specific order:

☐ First, when prompted for the standards data, navigate to and select the tab-delimited text file containing the consolidated IODC values for all standards.

☐ Second, when prompted for the sample data, navigate to and select the tab-delimited text file containing the IODs values for the specific specimen you are analyzing.

☐ Third, it will prompt for choosing the size of the x axis where you can change based on how big your genome size histogram is. Upon successful execution, the script will generate a histogram visualizing the genome size distribution for the analyzed specimen, see example in Figure 10.

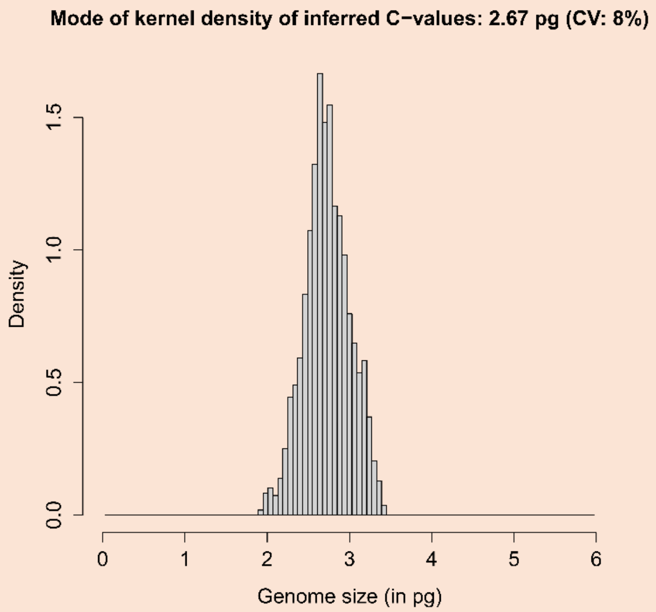


**Figure 10**
