## Supplementary File C. Codes and Excel templates for "Refining Feulgen: low-cost and accurate genome size measurements for everyone": 2. LM_model.R

```
library(Hmisc)
library(ggplot2)

Start<- read.table(file.choose(), header=T, sep="\t")

Data=Start[1:length(Start[,1]),]
 for (j in 1: length(Data[,1]))
+ {Data[j,]=as.numeric(Data[j,])}
reg=lm(Area~ X1.OD, data = Data)
summary(reg)
x=(Start$Area)
y=(Start$X1.OD)
library(Hmisc)
plot(x, y,
     xlab ="Nuclear area", ylab = "1/OD",
     pch = 19, frame =TRUE ,ylim = c(0,max(y+0.019)),xlim = c(0,max(x+1500)),xaxs = "i", yaxs = "i", col="#3399FF", minor.tick(nx = 3, ny = 3, tick.ratio = 0.5), cex=1.5, cex.lab=1.5, cex.axis=1.5, cex.main=1.5, cex.sub=1.5)

abline(lm(y ~ x, data = Start), col = "black", lwd = 1.7)
text(x = min(x+8000), y = max(y+0.009),label= paste("R�=",summary(reg)$adj.r.squared), col="black", cex=1.5)
text(x=min(x+17700),y=max(y-0.015), label=paste("Intercept p-value:",summary(reg)$coefficients[1,4]),col="black", cex=1.5)
```
