## Supplementary File C. Codes and Excel templates for "Refining Feulgen: low-cost and accurate genome size measurements for everyone": 4. LM_known_C-value.R

```
# --- 1. Load Libraries ---
library(ggplot2)
library(dplyr)
library(tcltk) 
# --- 2. Load Data ---
message("Select the TAB-separated file containing knowCv, IODaver, and Species columns")
Start_data <- read.table(tk_choose.files(caption = "Select TAB-separated data file"),
                         header = TRUE, sep = "\t", check.names = FALSE)

# --- 3. Check, Convert, and Prepare Data ---
required_cols <- c("knowCv", "IODaver", "Species") # Assuming these column names exist in your txt file
if (!all(required_cols %in% names(Start_data))) {
  stop("Error: Input file must contain columns named 'knowCv', 'IODaver', and 'Species'. Please check file.")
}

# Convert relevant columns to numeric
Start_data$knowCv   <- suppressWarnings(as.numeric(as.character(Start_data$knowCv)))
Start_data$IODaver <- suppressWarnings(as.numeric(as.character(Start_data$IODaver)))

# Convert the Species column to a factor
Start_data$Species <- as.factor(Start_data$Species)

# Remove rows with NA in plotting columns to prevent failure of the follwoing steps
plot_data <- Start_data %>% filter(!is.na(knowCv), !is.na(IODaver), !is.na(Species))

if(nrow(plot_data) == 0) {
  stop("Error: No valid data found in 'knowCv', 'IODaver', and 'Species' columns after conversion/filtering.")
}


# --- 4. Prepare Plot Parameters ---
min_x_val <- 0
max_x_val <- max(plot_data$knowCv, na.rm = TRUE)
min_y_val <- 0
max_y_val <- max(plot_data$IODaver, na.rm = TRUE)
base_text_size <- 11

# Make sure base_text_size is defined
base_text_size <- 11

# --- 5. Create the ggplot ---
# Ensure plot_data exists and has the required columns
if(exists("plot_data") && nrow(plot_data) > 0 && all(c("knowCv", "IODaver", "Species") %in% names(plot_data))) {

  gg_plot <- ggplot(plot_data, aes(x = knowCv, y = IODaver)) +

    # Add data points, colored by the Species factor
    geom_point(aes(color = Species), shape = 19, size = 4.5) +

    # Add regression line
    geom_smooth(method = "lm", formula = y ~ x, color = "black", se = FALSE, linewidth = 1) +

    # Define custom colors & labels for the levels "1" and "2"
    scale_color_manual(
      name = "Legend", # Updated Legend title
      # Values map colors to factor levels (ensure these match your data levels "1", "2")
      values = c("2" = "#e66101", # Color for P. americana (Level "2")  
                 "1" = "#fdb863"),  # Color for L. niger (Level "1")
      # Labels provide the display text, in order of factor levels ("1", then "2")
      labels = c("1" = expression(italic("L. niger")),
                 "2" = expression(italic("P. americana")))  
      ) +

    # Set labs and title
    labs(
      x = "Known C-values of standards",
      y = "Average IOD",
      title = "",
#IOD versus Known C-values (Overall Regression)
      color = "Species" # Define legend title
    ) +

    # Set axis limits starting from zero
    coord_cartesian(
      xlim = c(min_x_val, max_x_val + 0.5),
      ylim = c(min_y_val, max_y_val + 250000),
      expand = FALSE
      ) +

    # Use grid theme and apply text size adjustments
    theme_grey() +
    theme(
        axis.title = element_text(size = base_text_size * 1.5),
        axis.text = element_text(size = base_text_size * 1.5),
        plot.title = element_text(size = base_text_size * 1.5, hjust=0.5),
        legend.title = element_text(size = base_text_size * 1.3),
        # Use expression() for italicized labels in the legend text as well
        legend.text = element_text(size = base_text_size * 1.3),
        legend.text.align = 0,
        # --- Position legend inside plot: Bottom-Right ---
        legend.position = c(0.95, 0.1), # x=0.95 (near right), y=0.1 (near bottom) - Adjust
        legend.justification = c("right", "bottom"), # Align bottom-RIGHT corner of legend to position
        legend.background = element_rect(fill = alpha("white", 0.5)) # Optional: semi-transparent background
      )

  # --- 6. Print the Plot ---
  print(gg_plot)

} else {
   message("Error: 'plot_data' not found or has issues. Check previous steps.")
}
```
