## Supplementary File C. Codes and Excel templates for "Refining Feulgen: low-cost and accurate genome size measurements for everyone": 3. Genome_size_histogram.R

```
library(tcltk)
STANDARDS <- read.csv(tk_choose.files(caption = "Select a one-column file with DOI/C ratios of nuclei of one or several standards"))
SAMPLES <- read.csv(tk_choose.files(caption = "Select a one-column file with DOIs of nuclei of one sample for which you want to estimate the C-value"))

nb_standards=length(STANDARDS[,1])
nb_samples=length(SAMPLES[,1])

t=0
Cvalues=rep(0,nb_standards*nb_samples)
for (i in 1:nb_standards){
  for (j in 1:nb_samples){
    t=t+1        
    Cvalues[t]=SAMPLES[j,1]/STANDARDS[i,1]
  }
}

if (max(Cvalues) < 0.1) max_histo<-0.1
if (max(Cvalues) > 0.1) max_histo<-1
if (max(Cvalues) > 1) max_histo<-10
if (max(Cvalues) > 10) max_histo<-100

density_estimate<-density(Cvalues)
mode_value <- density_estimate$x[which.max(density_estimate$y)]
mode3SD <-signif(mode_value,3)

library(svDialogs)
max_histo <- dlgInput("Please enter below the maximum number of X axis to plot (or click OK to use the proposed default value):", default = max_histo)$res
max_histo <- as.numeric(max_histo)
left_density <- list("x" = density_estimate$x[1:which.max(density_estimate$y)], "y" = density_estimate$y[1:which.max(density_estimate$y)])
right_density <- list("x" = density_estimate$x[which.max(density_estimate$y):which.max(density_estimate$x)], "y" = density_estimate$y[which.max(density_estimate$y):which.max(density_estimate$x)])
cutoff <- 0.882*max(density_estimate$y)
left_cutoff <- left_density$x[which.min((left_density$y-cutoff)^2)]
right_cutoff <- right_density$x[which.min((right_density$y-cutoff)^2)]
sigma <- right_cutoff - left_cutoff
CV <- sigma/mode_value
CVpercent <- paste0(round(CV*100,0),"%)")

breakvalues=seq(max_histo*0.005,max_histo*0.995,by=max_histo/100)
histogram1 <-hist(Cvalues[Cvalues >= max_histo*0.005 & Cvalues < max_histo*0.995], xlab="Genome size (in pg)",ylab="Density", main=paste("Mode of kernel density of inferred C-values:", prettyNum(mode3SD),"pg (CV:", CVpercent),xlim=c(0,max_histo), prob=TRUE, breaks=breakvalues, cex.lab=1.3, cex.axis=1.3, cex.main=1.3, cex.sub=1.3, col="#99CCFF")

#col="#FF3300","#99CCFF", "#33FF00" 
lines(density_estimate, col = "red")
```
